# TMEM106B coding variant is protective and deletion detrimental in a mouse model of tauopathy

**DOI:** 10.1101/2023.03.23.533978

**Authors:** George A. Edwards, Caleb A. Wood, Quynh Nguyen, Peter J. Kim, Ruben Gomez-Gutierrez, Kyung-Won Park, Cody Zurhellen, Ismael Al-Ramahi, Joanna L. Jankowsky

## Abstract

TMEM106B is a risk modifier for a growing list of age-associated dementias including Alzheimer’s and frontotemporal dementia, yet its function remains elusive. Two key questions that emerge from past work are whether the conservative T185S coding variant found in the minor haplotype contributes to protection, and whether the presence of TMEM106B is helpful or harmful in the context of disease. Here we address both issues while extending the testbed for study of TMEM106B from models of TDP to tauopathy. We show that TMEM106B deletion accelerates cognitive decline, hindlimb paralysis, neuropathology, and neurodegeneration. TMEM106B deletion also increases transcriptional overlap with human AD, making it a better model of disease than tau alone. In contrast, the coding variant protects against tau-associated cognitive decline, neurodegeneration, and paralysis without affecting tau pathology. Our findings show that the coding variant contributes to neuroprotection and suggest that TMEM106B is a critical safeguard against tau aggregation.

## Introduction

TMEM106B was identified more than a decade ago as a risk modifier for frontotemporal lobar degeneration (FTLD) with TDP-43 inclusions (Van Deerlin et al., 2010). Since that discovery, SNPs associated with FTLD have been linked to elevated risk for multiple other neurological diseases including AD, LATE, and hippocampal sclerosis of aging (Nicholson and Rademakers, 2016; Feng et al., 2021). Even in healthy individuals, TMEM106B SNPs have been associated with transcriptional signatures of brain aging (Rhinn and Abeliovich, 2017). Given this growing association with disease risk, it may be surprising that the non-risk variants in the same untranslated region have been linked to cognitive resilience and neuroprotection. De Jager and colleagues identified TMEM106B as one of several genes contributing to cognitive preservation in subjects later found to harbor neuropathology normally associated with dementia (White et al., 2017). Subjects with these protective SNPs also appear to maintain a greater proportion of neurons in their brains in the face of aging and AD (Li et al., 2020).

Part of the difficulty in interpreting the role of individual SNPs in disease risk is that the entire TMEM106B gene is inherited in a single linkage disequilibrium block. Non-coding SNPs in the common (major) allele are associated with disease risk and segregate with the lone coding SNP that expresses threonine at position 185 (residue 186 in mice) (Van Deerlin et al., 2010; van der Zee et al., 2011; Gallagher et al., 2017). In contrast, complementary SNPs in the minor allele segregate with a coding variant that encodes a biochemically similar serine at this site (Nicholson and Rademakers, 2016). This highly conservative amino acid change would generally be discounted as having any influence on protein function or disease risk.

Nevertheless, two studies in transfected cells suggest that serine at this site may be active and protective (Nicholson et al., 2013; Jun et al., 2015). One study suggested that the serine variant decreased TMEM106B half-life to affect function, while the other suggested that it preserves autophagic flux relative to the threonine variant. Other studies disagree. Genomic analyses instead suggest that a non-coding SNP conveys disease risk vs resilience by changing chromatin structure to control TMEM106B expression levels (Gallagher et al., 2017). Only one study to date has tested the coding variant in vivo with disappointing results. The T186S variant had little impact on pathology or microgliosis with only mild changes in lysosomal enzyme activity and lipid content in a GRN deletion model of FTLD (Cabron et al., 2023). Despite the known association of the protective TMEM106B haplotype and its T185S coding variant with cognitive resilience and neuroprotection, no studies to date have examined these downstream consequences of disease in an experimental model.

What clues exist from prior studies suggest that reducing TMEM106B levels might be protective in disease. First, the original association study between TMEM106B and FTLD-TDP found that TMEM106B mRNA levels were greater in cases than controls and nearly three-fold higher in homozygotes for the major (risk) allele than the minor (protective) allele (Van Deerlin et al., 2010). Second, TMEM106B protein encoding S185 found in the protective allele has lower steady-state expression and a shorter half-life in vitro than the risk T185 variant (Nicholson et al., 2013). Third, the protective allele alters chromatin architecture in a way that is predicted to lower transcription compared with the risk allele (Gallagher et al., 2017). Fourth, incremental elevation of TMEM106B protein levels produces a corresponding rise in cytotoxicity (Gallagher et al., 2017).

Finally, aberrant TMEM106B fibrils have recently been identified in multiple disease settings where TDP-43 pathology was expected, suggesting a pathological build-up of a TMEM106B protein fragment with age and disease (Chang et al., 2022; Jiang et al., 2022; Schweighauser et al., 2022). Contradicting this seemingly strong support for protein reduction, in the four studies where TMEM106B deletion has been tested in GRN deletion models of FTLD-TDP, three show profound worsening of disease phenotypes (Klein et al., 2017; Feng et al., 2020; Werner et al., 2020; Zhou et al., 2020a). There is a growing appreciation of the impact TMEM106B has on disease risk, but no clear mechanism for how it does so.

Despite TMEM106B’s expanding association with a broad range of neurological diseases, surprisingly few studies have examined the impact of TMEM106B deletion in models other than FTLD-TDP. Further, only one prior study has examined the impact of the T186S coding variant, but with the same GRN deletion model of FTLD-TDP used to test TMEM106B deletion. Here we expand from FTLD-TDP to test whether TMEM106B deletion or coding variation are harmful or helpful in a mouse model of tauopathy. We use a well-studied MAPT P301S model to examine for the first time how genetic manipulation of TMEM106B affects cognitive decline, CNS degeneration, and transcriptional signatures of dementia. Our findings provide additional compelling evidence for a protective role of TMEM106B in models of neurodegenerative disease and extend that role from FTLD-TDP to the larger group of tauopathies. Moreover, our study offers the first evidence that the T186S coding variant is biologically active and neuroprotective in vivo.

## Results

### TMEM106B deletion worsens cognitive impairment and accelerates paralysis caused by pathogenic tau

Because so many of the diseases influenced by TMEM106B are associated with dementia, we began by examining how the presence or absence of TMEM106B influenced cognitive function in the context of tauopathy. We generated TMEM106B null mice using embryonic stem cells from the KnockOut Mouse Project (KOMP), which were crossed with ubiquitous Cre and Flp recombinase lines to remove the lacZ cassette and delete the essential exon 4. Unlike past work with KOMP mice in which both the lacZ and exon remained intact (Klein et al., 2017; Nicholson et al., 2018), our strategy produced nearly complete loss of TMEM106B protein. The TMEM106B deletion mice were intercrossed with the PS19 model of tauopathy carrying the P301S MAPT mutation to generate four genotypes for study (WT, KO, tau, and KO:tau). Behavioral testing began at 8 mo of age with several assays of locomotor function. Animals that showed evidence of hindlimb paralysis were removed from further testing. Counterintuitively, healthy KO:tau animals showed hyperactivity in the open field arena (Fig S1A-B) but swam at the same speed and efficiency as other genotypes in the straight swim test, which supported further testing in water-based cognitive tasks (Fig. S1C-D).

We performed an extensive cognitive testing battery starting with Morris water maze to assess spatial reference memory (Fig. S2A-C), followed by repeated reversal learning (RRL) to assess cognitive flexibility (Fig S2D-E), and finally ending with radial arm water maze to assess spatial working memory (Fig S2F-G).

This battery revealed significant impairments across multiple tasks for both tau genotypes relative to WT. The KO:tau animals tended to be the poorest performing genotype of the four, but never distinct enough from tau to be significant.

We initially opted for behavioral testing at 8 mo to maximize our chances of seeing cognitive impairment in the tau animals compared to WT (Takeuchi et al., 2011; Min et al., 2015; Briggs et al., 2017). While aging animals to 8 mo, a subset of the tau animals developed hind-limb paralysis that required euthanasia, which is a known phenotype of the PS19 model (Yoshiyama et al., 2007). By recording the age of paralysis onset, we realized that TMEM106B deletion was accelerating the phenotype. KO:tau mice developed paralysis as early as 6 mo, while it was 8.6 mo before the first tau animal showed hindlimb weakness (Fig. 1A). This effect on paralysis was surprising given the locomotor hyperactivity seen in behaviorally tested KO:tau mice, nevertheless, the median age at euthanasia was 9.3 months for KO:tau vs 9.9 mo for tau. This led us to wonder if our behavioral experiments may have suffered from a survivorship bias, such that only the healthiest animals lived long enough to undergo cognitive testing.

**Figure 1.**
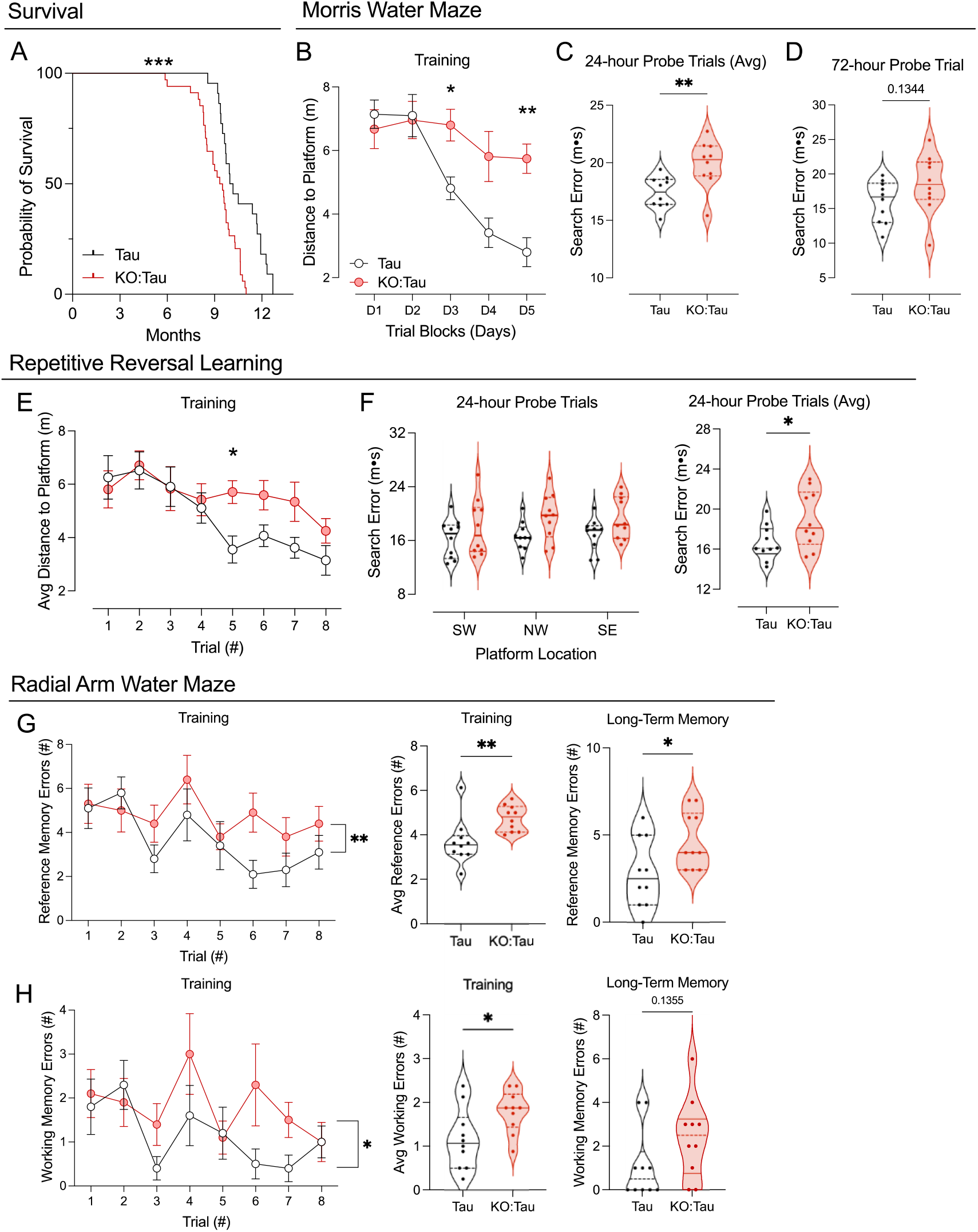
TMEM106B deletion accelerates paralysis and worsens cognitive behavior. A. Onset of hindlimb paralysis requiring euthanasia was measured for animals that were not euthanized after behavior. The age at paralysis onset was significantly earlier in KO:tau than tau mice. B. Spatial learning in the Morris water maze was assessed by the distance traveled to reach the escape platform. KO:tau mice performed significantly worse than tau animals on days 3 and 5 of training. C-D. KO:tau mice also recalled the escape location less well than tau animals during daily short term memory testing (C), but this difference did not persist during long-term memory testing (D). E. Cognitive flexibility measured by changing the platform location each day showed that KO:tau mice performed more poorly than tau mice on trial 5. Graph shows average performance across days as a function of trial number (8 trials/day). F. Short-term memory for the changing platform location was no different for any given location (left), but was worse overall in KO:tau mice when averaged across the three locations. G-H. During the learning phase of radial arm water maze, KO:tau mice made more incorrect arm entries (G, reference memory, left) and more repeated arm entries (H, working memory, right and middle graphs). Long-term memory was also impaired in KO:tau mice compared to tau when measured as incorrect arm entries (G, right), but did not reach significance for repeated arm errors (H, right). See Table S1 for ANOVA statistics. * p<0.05, ** p<0.01, *** p<0.001.

To overcome this survivorship bias, we examined a younger cohort of animals starting at 6 mo of age, when >90% of the KO:tau mice were still healthy. We focused these experiments on tau and KO:tau groups because we had seen no significant differences between WT and KO in our initial studies at 9 mo of age (Fig S2). We repeated the same 3 tasks in the same order as before, but with considerably different outcomes.

First, the KO:tau mice showed no hyperactivity or thigmotaxis in open field testing, suggesting that these phenotypes arose between 6 and 9 mo of age in the surviving KO:tau animals. As before, KO:tau mice swam as well as tau animals, both in the straight swim test and when evaluated in the cued platform trials following MWM (Fig S1E-F). Despite similar locomotor performance, cognitive outcomes at 6 mo were strikingly different than at 9 mo. In MWM, the KO:tau mice learned the hidden platform location less efficiently than tau animals swimming farther to the escape platform on days 3-5 of the 5 day training period (Fig 1B). KO:tau mice also remembered the trained location more poorly during short-term probe trials (Fig 1C), but failed to reach significance in the long-term probe trial (Fig 1D). In RRL, KO:tau mice learned each new platform location less efficiently than tau mice (Fig 1E). Average short-term memory for the new platform locations was also less precise in KO:tau than tau (Fig 1F). In RAWM, KO:tau mice learned less effectively than tau animals, making more working memory errors (repeated entries to the same arm) and more reference memory errors (any incorrect arm entry) while learning the task and recalling it later (Fig 1G-H). We conclude from these findings that TMEM106B accelerated cognitive decline in the tau mice and that the effect was easier to detect at an earlier age.

### Expression of T186S coding variant protects against cognitive decline in tau mice

The only common coding variant found in TMEM106B encodes a relatively conservative change from threonine to serine at position 185 in the human gene or 186 in the mouse. This variant is in linkage disequilibrium with multiple non-coding SNPs that associate with differential risk for various neurodegenerative disorders including FTLD and AD. Past work has disagreed on whether this coding variant plays any role in the protective effect of the minor allele, or if instead non-coding SNPs in the 3’ UTR act at the transcriptional level to influence TMEM106B function (Nicholson et al., 2013; Jun et al., 2015; Gallagher et al., 2017). We set out to directly test whether the T185S variant could influence disease pathogenesis by introducing the equivalent coding substitution in mice using CRISPR/Cas9. We confirmed the mouse T186S variant by Sanger sequencing of the full exon and high-resolution melt point analysis to check for off-target mutations.

Three lines were expanded for basic characterization and one was selected for further study. Because past work had suggested that the in vitro expression of the human T185S variant reduced TMEM106B protein half-life and steady-state levels, we tested TMEM106B expression in our T186S mice but found no differences from WT at any age examined (Fig S3) (Nicholson et al., 2013). T186S mice were intercrossed with PS19 tau to generate four genotypes for study (WT, KI, tau, and KI:tau); both KI and KI:tau mice were homozygous for the T186S variant. Animals underwent the same behavioral testing procedure as for KO x tau, starting at 8 mo of age.

Initial assessment in the open field and straight swim showed no difference between genotypes (Fig S4A-B), so we proceeded to the cognitive test battery. This is where the genotypes started to separate in surprising ways. Although we found no difference in learning the MWM escape location, there were pronounced differences in both short- and long-term memory for the trained location (Fig 2A-C, see Fig S5 for complete line graphs including all 4 genotypes). The preservation of short-term memory in KI:tau mice compared to tau alone reached significance in the MWM day 4 probe trial, and approached significance when averaged across days (p=0.06, Fig 2B). The effect became more pronounced for long-term memory, with KI:tau mice performing as well as WT (Fig 2C). Repeated reversal testing confirmed a protective effect of the KI allele, with KI:tau mice learning the new platform locations more efficiently and recalling them more accurately than tau animals (Fig 2D-E). Cognitive preservation was also observed in the RAWM, where KI:tau mice made fewer incorrect arm entries than tau animals during training and showed better memory for the escape arm during long-term probe tests (Fig 2F-G). Collectively, these results demonstrate that the serine variant imparts resilience against cognitive impairments caused by pathogenic tau.

**Figure 2.**
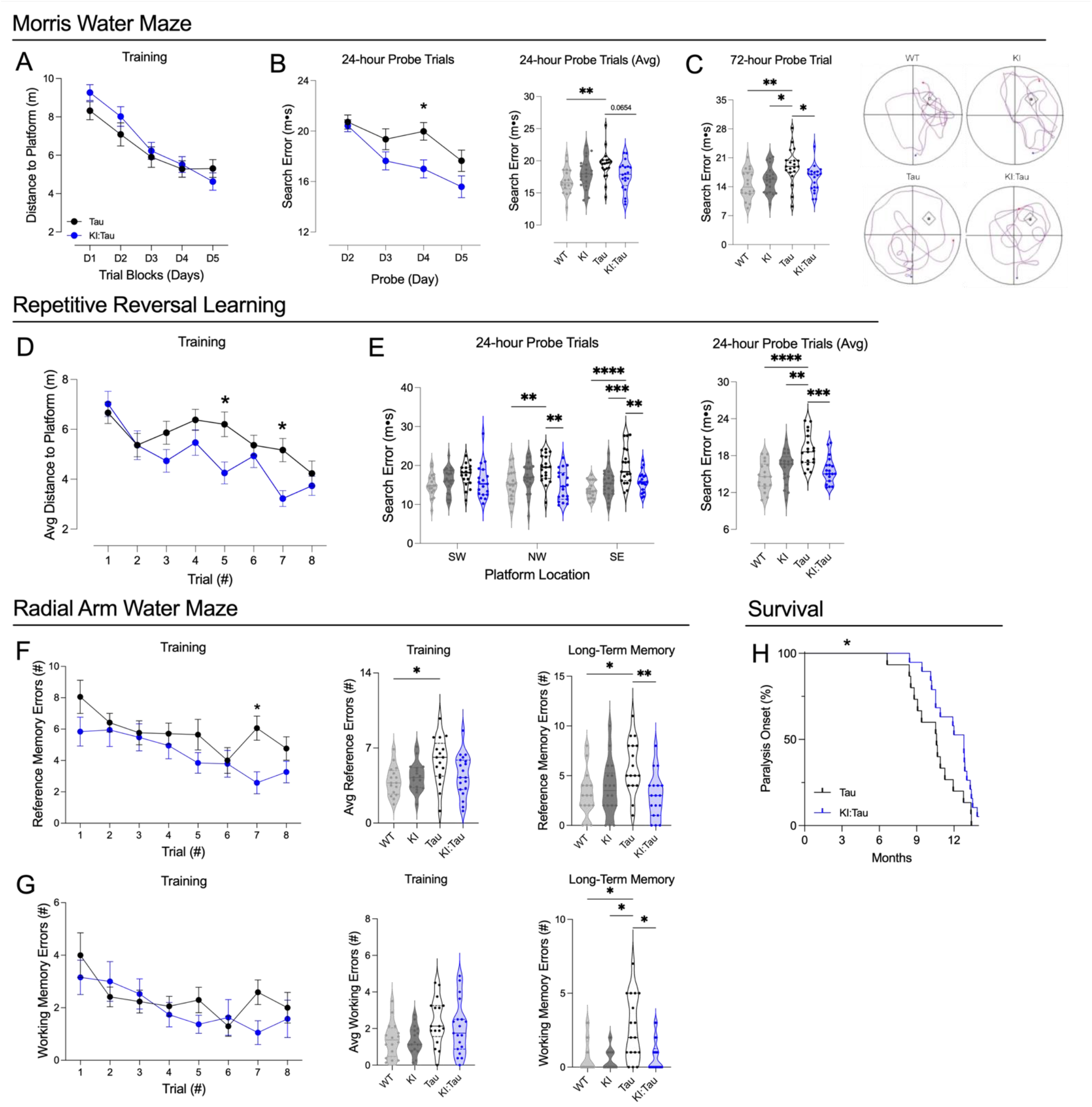
The T186S coding variant protects against cognitive decline and delays paralysis in tau mice. A. No difference was noted between KI:tau and tau groups during the learning phase of Morris water maze, measured as distance to the escape platform. B. Short term memory for the escape location was better in KI:tau mice than tau on day 4 (left). When short-term memory in daily probe trials was averaged across days, the improvement in KI:tau vs tau mice was nearly significant at p=0.065 (right). C. Long-term memory for the escape location was better in KI:tau mice than tau animals. Track plots show example search paths for one animal of each genotype during the long-term probe trial (right). D. Cognitive flexibility measured by changing the platform location each day showed that KI:tau mice performed better than tau mice on trials 5 and 7. Graph shows average performance across days as a function of the trial number (8 trials/day). E. Short term memory for each new platform location was better in KI:tau than tau mice by the second day of testing (platform location NW), and this difference widened with the third day of testing (platform location SE, left). On average, KI:tau mice showed better short-term memory for the changing platform locations than tau mice (right). F-G. During the learning phase of radial arm water maze, the number of incorrect arm entries made by KI:tau was lower than in tau mice on trial 7 (F, reference memory, right), but did not reach significance when averaged across trials (F, middle). Both genotypes made the same number of repeated arm entries during training (G, working memory, right and middle graphs). Remarkably, KI:tau mice recalled the trained escape arm better than tau animals during long-term testing, both when measured as incorrect arm entries (F, reference errors, right) and when measured as repeated arm entries (G, working errors, right). H. Onset of hindlimb paralysis requiring euthanasia occurred significantly later in KI:tau than tau mice. Significance values shown here reflect ANOVA testing with all four genotypes. * p<0.05, ** p<0.01, *** p<0.001, **** p<0.0001. See Table S1 for ANOVA statistics.

Because we had noted such a profound impact of TMEM106B deletion on the onset of paralysis in the PS19 mice, we extended this analysis to the KI colony. In contrast to the accelerated phenotype observed in KO:tau animals, expression of the T186S allele slowed the onset of hind limb paralysis from a median survival of 9.2 mo in the tau animals to 10.7 mo in KI:tau mice (Fig 2H). The minor coding substitution again had a pronounced impact on neurobehavioral phenotypes in this model.

### Loss of TMEM106B enhances neurodegeneration and pathologic markers of tauopathy

Our next challenge was to understand if and how each of our TMEM106B manipulations affected neuropathology to evoke such profound differences in cognitive function. Starting with the KO, we considered multiple mechanisms by which loss of TMEM106B may have accelerated cognitive decline and paralysis, however the most obvious is by hastening tau aggregation and its downstream consequences. We examined this question in a subset of mice that were behaviorally tested at 8 mo, and began by asking whether the loss of TMEM106B exacerbated neurodegeneration in tau mice. The Campbell-Switzer silver stain provided good tissue contrast for this analysis and allowed us to measure ventricular, hippocampal, and cortical volume in the same sections. We found that deleting TMEM106B in tau mice profoundly worsened neurodegeneration by all three measures (Fig 3A-B). Total ventricular area was 2.3 times greater in KO:tau than tau animals (Fig 3B). Hippocampal and cortical volumes were both decreased in KO:tau relative to tau. Hippocampal volume in KO:tau was ∼30% smaller than in tau mice; cortical volume was reduced by 10%.

**Figure 3.**
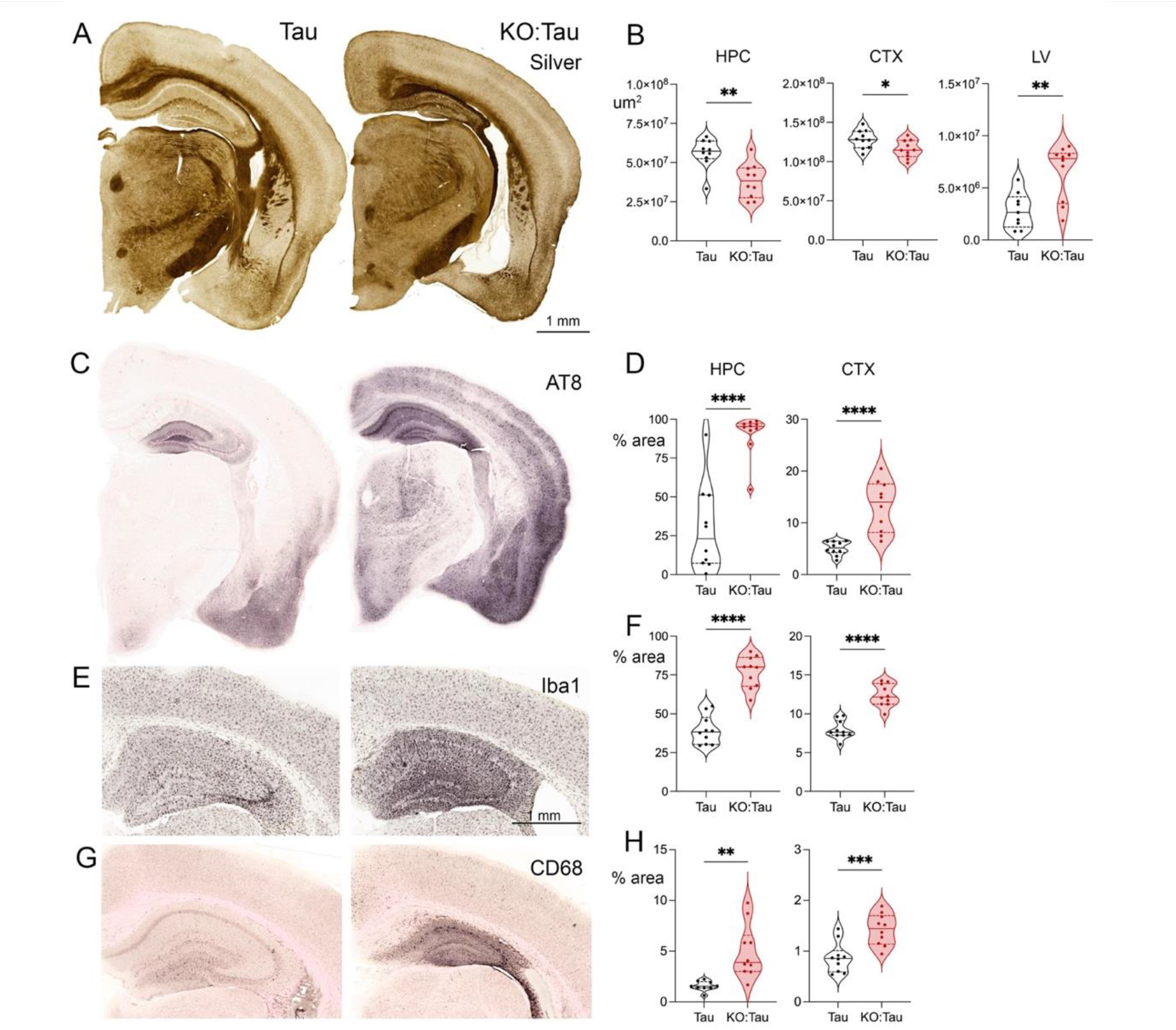
TMEM106B deletion worsens degeneration and tau pathology. A. Campbell-Switzer silver staining did not identify tangles, but did reveal marked ventricular enlargement in the KO:tau animals compare to tau alone. B. Silver-stained sections were used to measure the summed area of sections spanning the hippocampus (HPC), overlying cortex (CTX) and adjacent lateral ventricle (LV) in the tau and KO:tau mice to demonstrate a quantitative difference in the volume of all three regions upon TMEM106B deletion. C-H. Immunostaining and quantitation of AT8 (C, D), Iba1 (E, F), and CD68 (G, H) all demonstrate marked worsening of tau pathology and microgliosis upon TMEM106B deletion. Graphs in D, F, G show the percent area stained by each marker within hippocampus (HPC) and cortex (CTX). * p<0.05, ** p<0.01, *** p<0.001, **** p<0.0001

Tau pathology was also worsened by loss of TMEM106B. Phospho-tau AT8 staining occupied roughly 30% of the hippocampus in tau mice, but more than three times this area in KO:tau animals (Fig 3C, D). Neurofibrillary tangles identified by Gallyas staining were also more frequent in the KO:tau mice (data not shown), and appeared as early as 6 mo of age in both male and female KO:tau animals (Fig S6). Microgliosis identified by Iba1 staining tracked with pathological tau accumulation. Iba1 staining occupied ∼40% of the hippocampus in tau mice, but nearly twice that area in KO:tau mice (Fig 3E, F). CD68 immunostaining as a marker of phagocytic microglia also rose dramatically in KO:tau mice (Fig 3G-H). Overall, the loss of TMEM106B accelerated tau accumulation and worsened neurodegeneration resulting from tauopathy.

### TMEM106B T186S protects against degeneration without changing tau pathology

Given that the loss of TMEM106B so dramatically worsened both cognitive impairment and neuropathology, we predicted that the T186S variant which had preserved cognitive function would do so by diminishing neuropathology. Instead, we found that the coding variant protected against neurodegeneration without influencing phospho-tau pathology or microgliosis. Cortical volume was 13% larger in KI:tau mice and lateral ventricle ∼50% smaller than in tau animals, without changing hippocampal size (Fig 4A, B). Preservation of cortical volume was not accompanied by any difference in AT8 (Fig 4C-D), Iba1 (Fig 4E-F), or CD68 burden (Fig 4G-H). These findings suggest that the T186S variant protects against degeneration downstream of tau, but does not alter tau burden itself.

**Figure 4.**
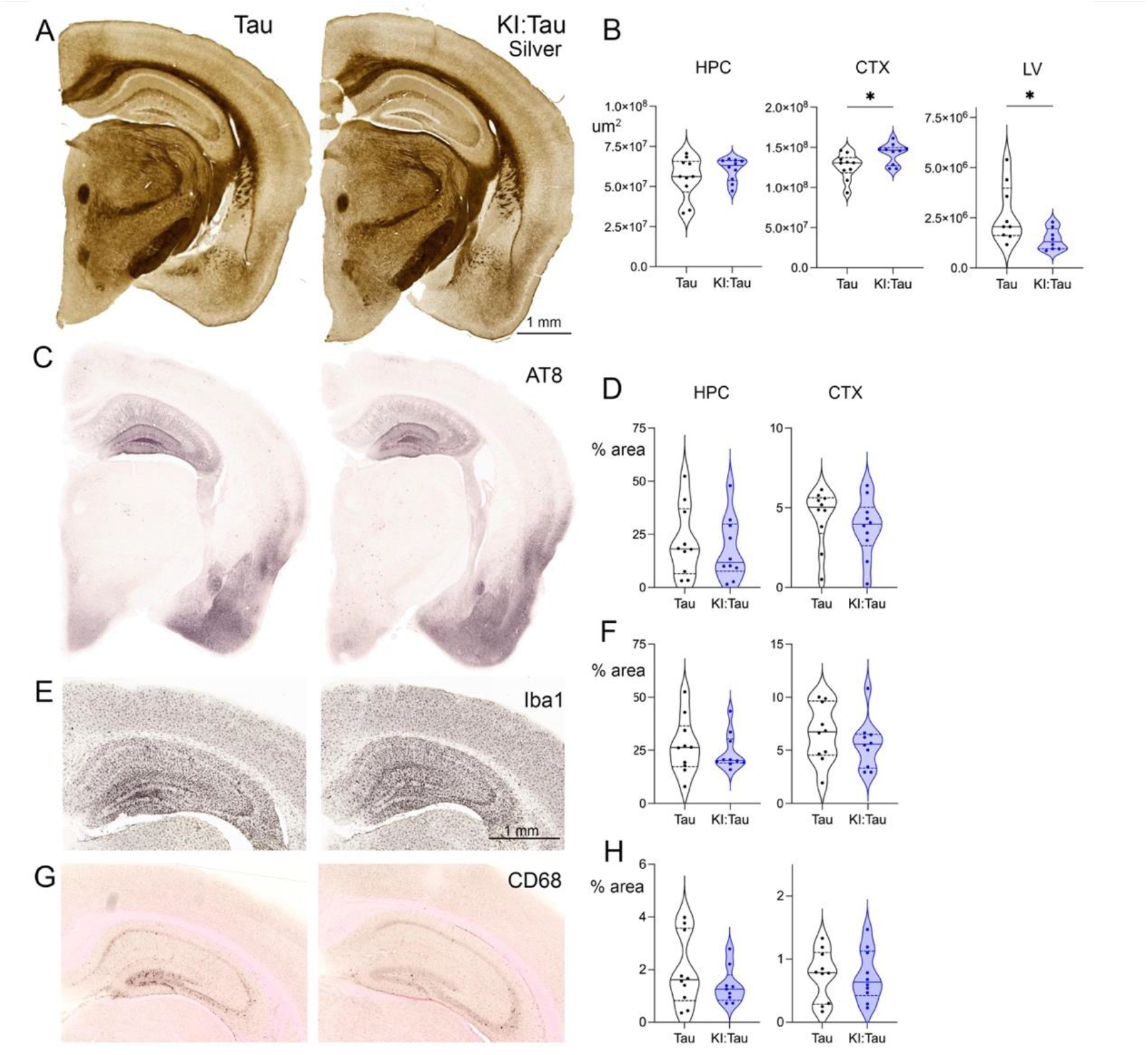
TMEM106B T186S protects against degeneration without changing tau pathology. A. Campbell-Switzer silver stain hinted at ventricular preservation in T186S KI:tau mice compared with tau alone. B. The summed area of hippocampus (HPC), cortex (CTX), and lateral ventricle (LV) were measured from silver-stained sections spanning the full extent of hippocampus. Tau animals carrying the T186S variant retained more cortical tissue and showed less ventricular enlargement than tau alone. C-H. Immunostaining and percent area quantitation of AT8 (C, D), Iba1 (E, F), and CD68 (G, H) were no different between genotypes in hippocampus or cortex. All images are from male mice. * p<0.05.

### TMEM106B deletion increases transcriptional overlap with AD

We next turned to RNA sequencing to see if a broad view of transcriptional changes between genotypes might provide clues about how TMEM106B protects from tauopathy. We performed mRNA sequencing on hemi-forebrain tissue from 5-7 behaviorally-tested mice of each genotype. We found few differences between WT and KO, in line with past studies on other TMEM106B deletion models ((Feng et al., 2020; Werner et al., 2020) but see (Zhou et al., 2020b), Table S2). We found far more differentially expressed genes (DEGs) between KO:tau and tau, where 170 genes were downregulated and 371 upregulated at an FDR cutoff of <0.05 (Table S2).

To gain functional insight into the DEGs distinguishing KO:tau from tau, we first performed a GSEA enrichment score (ES) analysis against the KEGG Alzheimer’s disease (AD) signature, which revealed significant enrichment of genes in this pathway (ES of 1.71, FDR q=0.06; Fig 5A). Surprisingly, testing against the entire KEGG database also revealed significant enrichment for genes common across Parkinson’s disease, Huntington’s disease, and AD modules, suggesting that loss of TMEM106B may be detrimental in multiple disease settings (Table S3). Next, we directly compared the KO:tau vs tau DEGs against multiple AMP-AD human brain transcriptional datasets (Bennett et al., 2012; Allen et al., 2016; De Jager et al., 2018; Mostafavi et al., 2018; Wang et al., 2018; Wan et al., 2020). A significant number of mouse DEGs overlapped and were concordant with human DEGs identified from comparison of AD vs healthy control (HC) postmortem brain tissue (134 upregulated, overlap Fisher p<0.0001 and 56 downregulated, overlap Fisher p=0.0004, Table S4). The correspondence between KO:tau and human AD was more clearly visualized by plotting the expression level of each tau genotype against the expression level of AMP-AD human AD for all DEGs identified from comparison of KO:tau vs tau. The resulting correlation between KO:tau and AD was significantly stronger than for tau and AD, and was consistent when plotting expression from human whole brain or specific regions such as temporal cortex, parahippocampal gyrus or superior temporal gyrus (Fig 5B and C).

**Figure 5.**
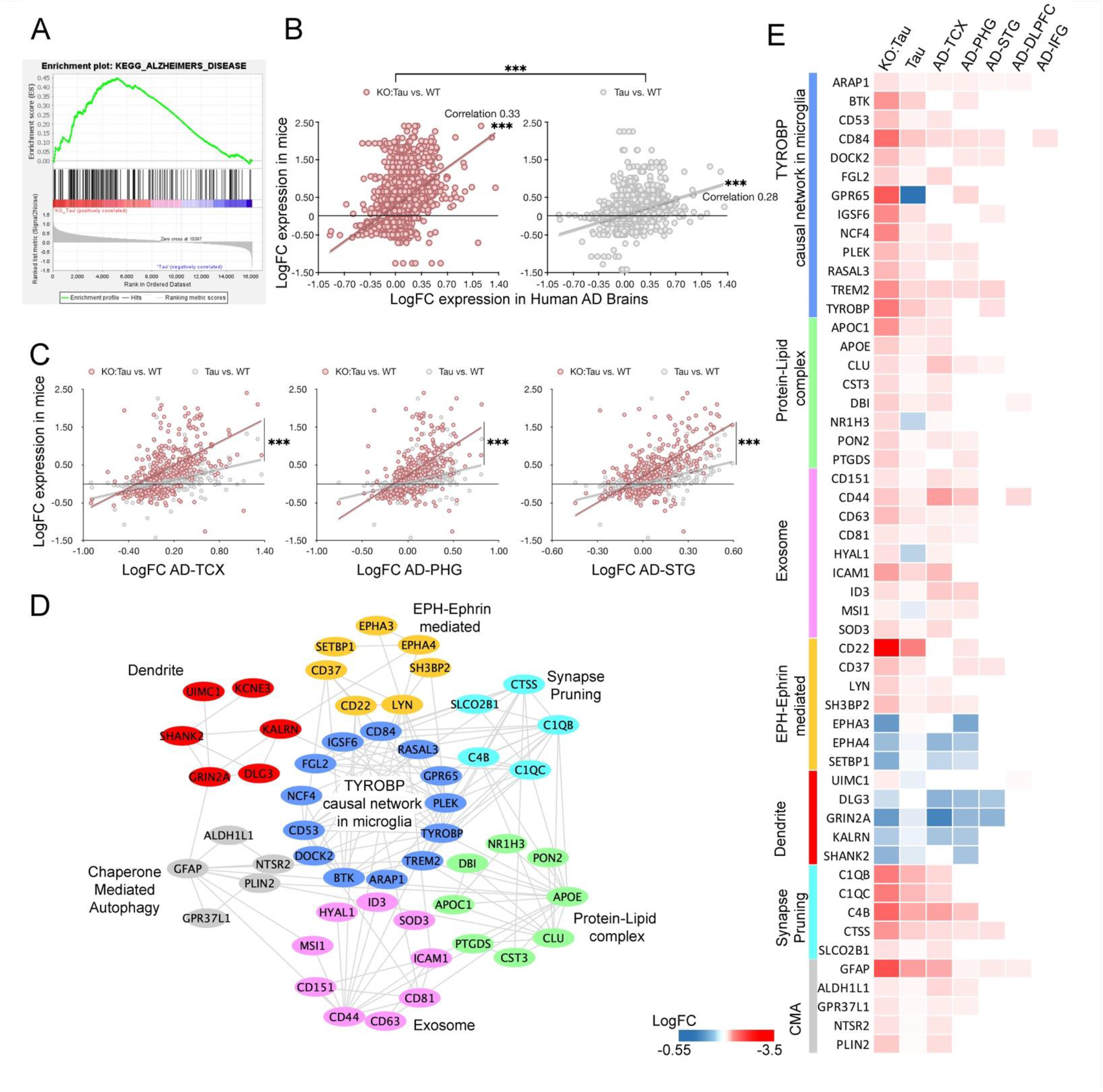
TMEM106B deletion improves the transcriptional alignment between tau mouse brain and human AD. **(A)** GSEA Enrichment Score (ES) analysis reveals enrichment of AD-KEGG pathway genes in KO:tau transcripts vs tau alone. **(B)** Relationship between human AD and mouse brain for DEGs identified by comparison of KO:tau vs tau. Values for human AD gene expression (relative to HC) were derived from the AMP-AD database for 5 human AD brain regions on the X axis (log2 fold change) against the corresponding expression level for KO:tau (left panel) or tau (right panel; both relative to WT). Correlation with human AD expression is significantly higher in the KO:tau animals than in mice expressing tau alone. **(C)** Same as B, but showing values for each human brain region separately (pink=KO:tau vs WT, grey=tau vs WT; TCX=temporal cortex, PHG=parahippocampal gyrus and STG=superior temporal gyrus). **(D)** Network clustering showing several enriched modules among the KO:tau vs tau DEGs that overlap and change in the same direction as human AD brain. Colors distinguish separate, but related, functional modules. **(E)** Heat map illustrating DEGs used to generate functional modules in panel D, plotted to show the degree of change against WT (mouse) or HC (human) controls. TMEM106B deletion increases the magnitude of change towards a more AD-like expression pattern compared to tau alone. DEG expression is shown for both KO:tau and tau groups against 5 different human AD brain regions (DLPFC=dorsolateral prefrontal cortex, IFG=inferior frontal gyrus). *** p<0.001.

To gain insight into the specific mechanisms being affected by loss of TMEM106B, we assessed the enrichment of functional transcription signatures in KO:tau vs tau mice using STRINGdb. This analysis revealed enrichment in the TYROBP causal network associated with microglia, which also included synapse pruning genes (Table S5, and Fig S7). Other pathways related to microglia and inflammation included chemokine signaling, cytokine-cytokine receptor interaction, and microglia phagocytosis. In line with TMEM106B function in autophagy and endo-lysosomal trafficking, we observed enrichment in pathways related to lysosomal lumen peptidases and hydrolases, chaperone mediated autophagy, and exosome/secretory granules. There was also a high degree of overlap and concordance with human AMP-AD transcriptional signatures and functional pathways (Fig 5D-E, Table S6). Restated, the signature of TMEM106B deletion in tau mice paralleled that of human AD. Given the overlap of microglial signatures from KO:tau and AD, we next tested their correlation with disease-associated microglial genes (DAMs) and plaque-induced genes (PIGs) indicative of microglial activation in disease models (Keren-Shaul et al., 2017; Chen et al., 2020). By measuring expression differences for each tau genotype against WT and then plotting those values against the corresponding value in human AD, we again found that TMEM106B deletion increased the correlation between mouse tau and human AD both overall and within several of the individual brain regions tested (Fig S8). Taken together, this data suggests that loss of TMEM106B pushes PS19 tau mice towards a transcriptional profile that better represents human AD than tau alone.

### TMEM106B T186S elicits only mild transcriptional changes in the tau brain, but hints at an anti-inflammatory mechanism

We next performed similar transcriptomic analysis with the KI:tau mice. There were surprisingly few DEGs that met an FDR cutoff of 0.05 (23 genes, Table S2). We therefore relaxed the stringency to see if this revealed any functional insights (p<0.01, Table S2). Since we observed amelioration of behavioral decline, paralysis, and neurodegeneration in KI:tau mice, we expected to see a decrease in transcriptional correlation between KI:tau and human AMP-AD compared with tau alone. Surprisingly, that was not the case. The DEGs identified from KI:tau vs tau at p<0.01 were positively correlated with human AD for KI:tau mice, however the same genes were not correlated between mouse and human for tau alone (Fig 6A-B). Prompted by this, we next asked whether we might gain insight into the molecular mechanisms at play in the KI mice by analyzing DEGs that distinguished KI:tau from tau. Few DEGs from the comparison of KI:tau vs tau overlapped with DEGs from the corresponding KO comparison, suggesting that TMEM106B deletion and the coding variant elicit distinct molecular changes in tauopathy (31 overlapping out of 282 KI:tau and 1370 KO:tau DEGs at p<0.01). Since KO:tau DEGs enriched for modules that promote disease, while our behavior and neurodegeneration analyses showed that the coding variant attenuated progression, we hypothesized that DEGs in KI:tau mice may represent compensatory transcriptional changes in AD that protect neurons from degeneration. Functional enrichment analysis suggested this may be partially true: pathway analysis of KI:tau DEGs identified two different modules consistent with attenuation of the inflammatory response (anti-inflammatory cytokine production and regulation of macrophage cytokine production, in addition to increased IL-10R expression (Larson et al., 2010; Ziogas et al., 2014; Davies et al., 2015; Sadagurski et al., 2015; Chen et al., 2016; Waqas et al., 2017; Burmeister and Marriott, 2018; Li et al., 2018; Heuberger and Schuepbach, 2019; Zhang et al., 2019; Cen et al., 2022) (Fig 6C and Fig S9; Tables S7 and S8). This data suggests that the coding variant may act by countering inflammation downstream of tauopathy to preserve neuronal integrity and cognitive function.

**Figure 6.**
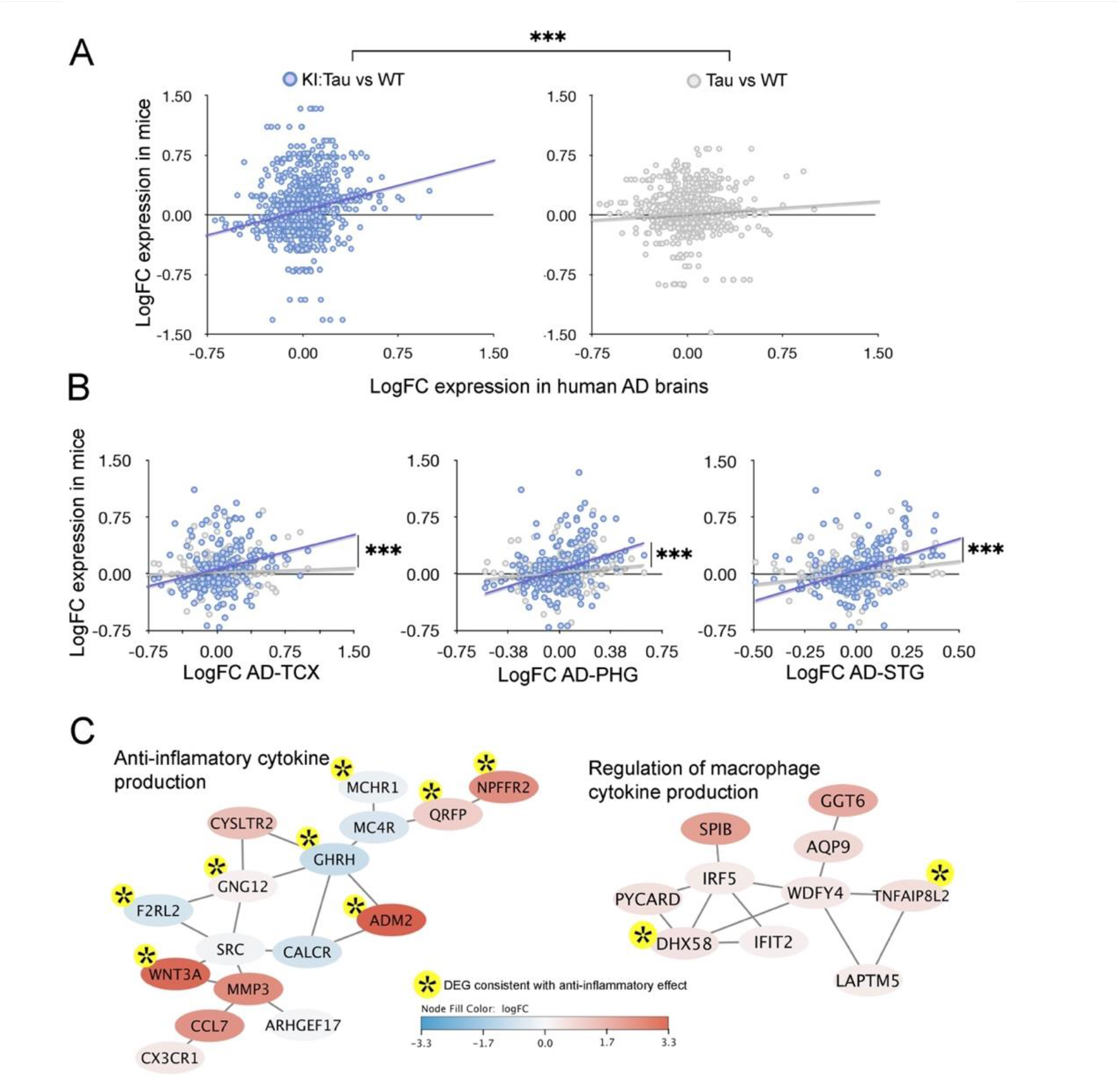
Transcriptomic analysis of KI:tau brain reveals presumptive compensatory transcriptional modules. A. Mouse brain expression as a function of fold change for human AD (KI:tau vs WT and tau vs WT against AD vs HC, log2 fold change). Genes plotted here are DEGs identified from comparison of KI:tau vs tau at p<0.01. KI:tau was positively correlated with AD, while tau alone was not for this gene set. B. Same as A, but plotted separately for individual brain regions. TCX = temporal cortex, PHG = parahippocampal gyrus, STG = superior temporal gyrus. C. Networks showing two functionally enriched modules among DEGs derived from comparison of KI:tau vs tau brain signatures. Both modules overlap with human AD, including DEGs that change in the same direction for mouse and human. Node color indicates the direction and magnitude of transcriptional change in KI:tau mice. Asterisks highlight nodes where the transcriptional change is consistent with an attenuation of the inflammatory response. *** p<0.001

## Discussion

Past work to understand the role of TMEM106B in the brain has focused on mouse models of FTLD-TDP due to the early, repeated, and strong GWAS associations with this subtype of FTD and the GRN mutation in particular (Nicholson and Rademakers, 2016). Yet since those landmark GWAS studies in FTLD-TDP, TMEM106B has been shown to modify risk for a variety of other neurological disorders including AD, CTE, cognitive decline in PD and ALS, LATE, and age-related hippocampal sclerosis, which all share cognitive decline as a key feature (Feng et al., 2021). More recently, TMEM106B fibrillar aggregates have been discovered in a range of dementias including FTLD-17, AD, ARTAG, LNT, and PSP that are characterized by tauopathy (Perneel and Rademakers, 2022). Here we provide the first experimental test of how genetic manipulation of TMEM106B affects disease progression in a mouse model of tauopathy, and the first test of its impact on tau-associated cognitive decline. Our work offers four main discoveries to the growing literature on this elusive protein. First, we show that endogenous TMEM106B is protective against tau pathology and tau-induced cognitive dysfunction. Loss of TMEM106B in MAPT P301S mice profoundly worsened every outcome we measured. Second, TMEM106B deletion significantly increased the transcriptional alignment between tau mice and human AD. To the best of our knowledge, this is the first time that TMEM106B manipulation has been shown to improve transcriptional overlap with human disease. Third, our findings demonstrate that the T186S coding variant in TMEM106B is biologically active and limits both neurodegeneration and cognitive decline due to tauopathy. Finally, most broadly, our study demonstrates that TMEM106B plays an important protective role in models of neurodegenerative proteinopathy beyond TDP-43.

Prior studies of TMEM106B deletion in the GRN null model of FTLD-TDP produced conflicting results with diametrically opposed outcomes. The first study to appear showed TMEM106B deletion to be protective against a subset of lysosomal abnormalities, locomotor hyperactivity, and behavioral disinhibition caused by GRN deletion, without altering microgliosis, lipofuscinosis, or retinal degeneration (Klein et al., 2017). Three subsequent studies in the same GRN model found instead that TMEM106B deletion either worsened existing abnormalities, or more often, synergized with GRN to evoke new deficits in lysosomal function, locomotor behavior, gliosis, neurodegeneration, and neuroinflammation (Feng et al., 2020; Werner et al., 2020; Zhou et al., 2020a). Our findings are better aligned with these three later studies in concluding that TMEM106B is normally protective against multiple aspects of neurodegeneration associated with proteinopathy. We show that TMEM106B deletion accelerated or exacerbated cognitive decline, hindlimb paralysis, phospho-tau and NFT accumulation, microgliosis, and neurodegeneration. Loss of TMEM106B had a marked effect on the brain transcriptional profile of tauopathy with a particular impact on signatures of microglial function. Collectively, we interpret this to suggest that endogenous TMEM106B acts upstream of tau pathology to slow aggregation and thereby delay downstream effects on behavior, gliosis, and degeneration. Surprisingly, this protective function is primarily uncovered in the context of cellular stress. Like others before us, we find only mild or no effect on locomotion and only a small number of transcriptional changes between WT and KO mice, (Klein et al., 2017; Feng et al., 2020; Luningschror et al., 2020; Werner et al., 2020) but see (Zhou et al., 2020b). Past work has shown that TMEM106B can influence myelination, lysosomal pH, size, and movement (Feng et al., 2021), but it is under the stress of aging and disease that this protein appears to have greatest impact.

Our transcriptional analysis of KO:tau mice confirmed and expanded our understanding of the broad impact TMEM106B deletion had on tau-associated pathological processes. TMEM106B deletion pushed the mouse tau transcriptional signature to better align with that of human AD, perhaps making it a better model of the disease than tau alone. Network modules enriched for mouse DEGs overlapped significantly with modules altered in human AD. These modules covered a range of functions from synapse pruning to chaperone-mediated autophagy that were highly connected to the TYROBP signaling pathway, consistent with the idea that TMEM106B plays a major role in the microglial response to neurological insult. While many microglial-associated pathways were exacerbated, several neuronal structure and functional pathways were downregulated, confounding any attribution of primary vs secondary cellular effects in promoting disease. The upregulation of exosome-related genes suggests a mechanism by which loss of TMEM106B might promote tau spreading, but without hinting at the cell type responsible. Nevertheless, the broad range of transcriptional modules impacted by TMEM106B deletion and the various cell types they represent support the idea that endogenous TMEM106B gates an early pathological event to slow disease.

Our work also uncovered a sharp contrast between disease acceleration caused by TMEM106B deletion and neuroprotection afforded by the T186S coding variant. First and foremost, our study is the first to demonstrate a biological effect of this variant in a disease model. This was unexpected because past work suggested the causal SNP was in the 3’UTR, where it altered chromatin architecture in a manner predicted to diminish TMEM106B transcription (Gallagher et al., 2017). Disease risk associated with the major allele was considered independent of the protein sequence. Where protein sequence was shown to have an effect, T185S was thought to exert disease protection by diminishing protein half-life. Here we show that the T186S variant protects against cognitive decline, hindlimb paralysis, and neurodegeneration, and does so without altering TMEM106B steady-state protein levels. We were surprised to find that the phenotypic changes observed in KI:tau mice were opposite those seen in KO:tau animals. The difference was particularly striking for the two functional outcomes of cognition and mobility. We were even more surprised to discover that this functional protection arose without any obvious change in tau pathology. This outcome is similar to the findings of a recent study testing the T186S variant in a GRN deletion model of FTLD-TDP (Cabron et al., 2023). T186S had no impact on lysosome and lipofuscin accumulation that are pathological hallmarks of GRN deletion, and showed only a limited effect on microgliosis that was restricted to CD68 elevation in the thalamus. This study didn’t report on retinal neurodegeneration or motor impairment that are also features of the GRN deletion model. Our discovery of cognitive, motor, and neuroprotection in the PS19 mice suggest that T186S may act downstream of tau aggregation here, and perhaps downstream of lysosomal dysfunction in the GRN model.

Given the pronounced behavioral and neuroprotective effects of the T186S variant - and the dramatic transcriptional changes we found with TMEM106B deletion - we were frankly disappointed at the minimal insight that could be gleaned from the transcriptional profile of KI:tau mice. Only 23 DEGs separated the KI:tau and tau groups using a standard FDR of 0.05. Even with a relaxed cutoff of p<0.01, few DEGs in the KI:tau vs tau comparison corresponded with the much larger set of genes separating KO:tau from tau, suggesting that the two TMEM106B manipulations affected disease phenotypes through distinct pathways. Unlike TMEM106B deletion, the coding variant had no impact on the transcriptional correlation between mouse tau and human AD. This limited transcriptional effect echoes similar futility with T186S in the GRN null animals (Cabron et al., 2023). Despite this drawback, we were able to mine DEGs using a relaxed cutoff for hints at what might underlie the variant’s protective effect. This analysis uncovered a weak anti-inflammatory signature in the KI:tau mice converging on cytokine production. Further studies are needed to confirm and explore this potential mechanism, beginning with the identity of cytokines affected by TMEM106B.

We recognize several limitations to the current study that should be considered when evaluating our results. First, our data contains significant inter-animal variability that arose from two main sources. The PS19 model was created on a mixed C3B6 genetic background which we maintained through intercross with TMEM106B KO and KI lines. This choice eased breeding with increased litter sizes and fewer litter losses, but also meant that no two animals had the same genetic background. The PS19 model is also known to have a pronounced sex difference with earlier pathological onset in males than in females (Zhang et al., 2012; Yanamandra et al., 2013). All of our experimental groups included both sexes. These two factors mean that we likely missed subtle differences between genotypes, but also that the significant differences we found are robust to sex and genetic variables. Another experimental limitation of our work is the use of intact forebrain tissue for transcriptional analyses rather than individual brain regions or single nuclei. This choice arose in part out of expedience during the COVID pandemic when we had limited time to collect tissue from many our initial animals. In retrospect, we might have better focused transcriptional analyses on dissected hippocampal tissue where pathological changes were most pronounced and where circuits responsible for spatial learning and memory are centered. As with the inter-animal variance, by including unaffected brain areas in our RNA sequencing, we likely missed small focal transcriptional differences between groups, but it also means that the differences we found were sizeable enough to emerge through substantial background noise.

Moving from GWAS to mechanism can be challenging under the best of circumstances and has been particularly fraught for TMEM106B. The very conservative coding substitution T185S yielded no obvious clue that it could affect protein function (Nicholson and Rademakers, 2016). TMEM106B lacks the smoking gun that can come from gene deletion in animal models as with TREM2 (Colonna, 2023), from protein structure for LILRB2 (van der Touw et al., 2017), or from known function as with CR1 (Zhu et al., 2015). Our understanding of TMEM106B has been further slowed by conflicting results in seemingly similar experimental models that hint at biology we have yet to untangle (Feng et al., 2021). While we have yet to identify the mechanism by which TMEM106B protects against tauopathy, or its coding variant against the downstream consequences of this pathology, we hope our results will bring the field a step closer to understanding this protein. We suspect that TMEM106B may play different roles depending on the cell type and the disease context. Future studies have much yet to discover.

## Methods

### Mice

#### TMEM106B deletion

Cryopreserved sperm for the line C57BL/6N-*Tmem106b*^tm2a(KOMP)Wtsi^ was purchased from KnockOut Mouse Project (KOMP) Repository at University of California Davis. This line was created from ES cell clone EPD0047_1_E02, generated by the Wellcome Trust Sanger Institute and made into mice by the KOMP Repository (www.komp.org) and the Mouse Biology Program (www.mousebiology.org) at the University of California Davis. The recovered sperm were used for in vitro fertilization of C57BL/6J oocytes by the BCM Genetically Engineered Mouse Core. 75 pups were recovered from 8 implanted females, of which 36 were heterozygous for the targeted allele. The resulting TMEM106B tm2a line was subsequently maintained by backcross to C57BL/6J. Tm2a mice were intercrossed with mice expressing Flp recombinase under control of the *β*-actin promoter which had been backcrossed to C57BL/6J for 8 generations (kind gift of Russell Ray, BCM, available from Jax as strain #3800 on a mixed background (Rodriguez et al., 2000)) to generate the conditional deletion tm2c line, which was again maintained by mating with C57BL/6J. Offspring from the tm2c line were mated with mice expressing Cre recombinase under control of the *β*-actin promoter which had been backcrossed to C57BL/6J for 3 generations (kind gift of Russell Ray, BCM, available from Jax as strain #19099 on a mixed background (Lewandoski et al., 1997)) to generate the final tm2d constitutive knockout allele lacking the targeted exon 4 and the floxed *β*-galactosidase/neomycin resistance cassette. The line was subsequently outcrossed with C3HeJ before being intercrossed with PS19 animals on a C3B6 background. Mice were genotyped by PCR using the following primers: Primer 58 (intron 3 upstream of first FRT site): TTC TCT CCA TGT GCT GCA TTA T, Primer 59 (just before exon 4 but inside the loxP sites; deleted by Cre): GAA TAC TTC TCT CCT TAG CCC TTT AG, Primer 60 (part of loxP site downstream of exon 4): GCG AGC TCA GAC CAT AAC TT. Animals homozygous for TMEM106B^tm2d^ will exhibit a band at 304 bp. Wild-type animals will exhibit a band at 496 bp, and heterozygous TMEM106B^tm2d^ will have bands at 304 bp and 496 bp.

*TMEM106B T186S knock-in*. The TMEM106B locus was targeted by CRISPR/Cas9 to change threonine 186 to serine within the endogenous mouse TMEM106B locus (homologous to human T185S). 200 C57BL/6J embryos were injected with 100 ng/µL Cas9 mRNA, 20 ng/µL guide RNA, and 100 ng/µL single stranded insert oligonucleotide by the BCM Genetically Engineered Mouse Core. The guide RNA sequence was: TTACCTGCTTCATATCAAGT, targeting PAM site at exon 6 of TMEM106b locus. A single stranded oligonucleotide was used as the donor template (AATAACTATTATTCTGTTGAAGTTGAAAACATCACTGCTCAAGTCCAGTTTTCAAAAACCGTGATTGGAAAGGCTCGTTTAAACAACATTTCGAACATTGGCCCATTAGATATGAAGCAGGTAAGCCTATTCAGGAGTTGAGGGG) to convert TMEM106B Thr 186 into Ser (ACT > TCG) with two additional silent mutations: I185I (ATA > ATT) was added to introduce a novel BstBI site, and L191L (CTT > TTA) was added to block re-cutting of the correctly edited genome. 209 injected embryos were transferred to pseudopregnant females, yielding 59 animals of which 35 were positive for the T186S allele. Three positive founders were expanded by mating with C57BL/6J wild-type animals. Offspring were screened by Sanger sequencing across the targeted region and by high resolution melting analysis for the top 4 predicted off-target sites; all sites except the intended target came back as wild-type. No differences in brain TMEM106B expression level were observed between the three knock-in lines and so further studies focused on a single line identified as 5825.

Sanger sequencing was repeated on tail DNA from 5825 offspring covering an expanded area surrounding the targeting site to include exon 6 plus approximately 50 intronic base pairs. The line was subsequently outcrossed with C3B6 before being intercrossed with PS19 animals on a C3B6 background. All offspring for study were generated from mating TMEM106B T186S heterozygous mice with PS19;TMEM106B T186S heterozygous mates. Animals were genotyped by PCR using the following primers: Primer 46: GTG TGT ATT TCC TGT CTC TGT T; Primer 47: GGG AGA GGA TGA GGG ATT T. All animals should exhibit a band at 407 bp before BstBI restriction digest at 65°C. T186S animals should exhibit a band at 97 bp and 208 bp WT should exhibit a band only at 407 bp, and heterozygous mice will have 3 bands at 97, 208, and 407 bp.

#### MAPT P301S transgenic line PS19

B6;C3-Tg(Prnp-MAPT*P301S)PS19Vle/J mice (hereafter referred to as Tau or PS19) were obtained from Jackson Laboratory (stock #8169) and maintained on a B6;C3 background. These mice express the 4R1N form of human MAPT encoding the P301S mutation under control of the mouse prion protein promoter as described by Yoshiyama et al. (Yoshiyama et al., 2007)

All animal experiments were reviewed and approved by the Baylor College of Medicine Institutional Care and Use Committee. Add anything about housing the mice in the satellite room during testing?

### Behavioral analysis

Animals began behavioral testing either at approximately 6 or 8 months of age. All animals were screened for paralysis prior to behavioral testing and eliminated from the study if present. Animals that developed hindlimb weakness during study were removed from further testing, but were included in tasks completed prior to the onset of paralysis. The number of animals from the 8 mo testing groups that were removed for paralysis at some point between open field (OF) and radial arm water maze (RAWM) was: tau = 2, KO:tau = 3; tau = 4, KI:tau =2. Additional mice from the 8 mo group could not be tested in RAWM due to the COVID-19 shutdown of our institute in March 2020, leaving group sizes for this task lower than for Morris water maze (MWM) or repeated reversal learning (RRL). Animals were included in analysis for all tasks they completed. Remaining animals underwent the complete behavioral battery at 8 mo: n= 18 WT, 18 KO, 21 tau, 20 KO:tau; n=18 WT, 21 KI, 22 tau, 21 KI:tau, divided roughly evenly between sexes. The 6 mo cohort included 10 tau and 10 KO:tau animals (4F/6M); all 6 mo animals completed the full behavioral battery. One KO:tau animal from the 8 mo cohort was subsequently removed from all analyses after it was discovered that this animal showed signs of visual impairment during visible platform testing.

Animals were handled for 5 min/d for 2 d prior to testing. Behavioral analysis began with open field (OF) on day 1, followed by straight swim on day 2, Morris water maze (MWM) on days 3-7 with final probe trial on day 10, followed by repeated reversal (RR) testing on days 10-12, visible platform water maze testing on day 13, and ending with RAWM on day 14 with final probe trial on day 17. Both male and female mice were used for testing in roughly equal numbers for all genotypes. All analyses were done using ANY-maze video tracking system (Version 4.99v, Stoelting Co.). Methods for OF, MWM, and RAWM are modified from Fowler et al. (Fowler et al., 2014). Animals were performed by a single experimenter blinded to genotype.

#### Open field

Testing was done in white acrylic open-top boxes (46 × 46 × 38 cm) in a room with indirect white fluorescent lighting to lower the brightness. Activity was recorded with an overhead camera for a single 30 min session.

#### Morris water maze

MWM testing was conducted in a circular tank measuring 58 cm high and 122 cm in diameter. The water level was 20 cm from the top of the tank and made opaque using nontoxic white paint. Water temperature was maintained between 19 to 21°C. As described in Fowler et al., before the start of acquisition training, mice received 1 d of training in a straight swim channel to acclimate them to the water and check for motor deficits. Mice received four trials with a 15 min intertrial interval (ITI) in a channel constructed of white acrylic and measuring 107 × 56 × 14 cm, which was placed in the center of the pool. Visible cues were removed from the room during straight swim shaping trials. Mice were allowed 60 s to reach a submerged platform on the opposite side of the channel. Mice that failed to reach the platform were guided to the location by the experimenter. Mice were allowed to stay on the platform for 15 s before being removed from the water, dried, and returned to their home cage placed on a heating pad between trials.

Acquisition training in the MWM consisted of four trials per day with a 15-20 min ITI. A square platform (10 × 10 cm) covered in nylon mesh for traction was located 1 cm below the surface in the NE quadrant of the maze, half-way between the side and the center of the pool. Mice were placed in the maze facing the wall at each of four randomized, cardinal start locations and allowed 60 s to locate the hidden platform utilizing visible cues around the arena. As with straight swim, animals that failed to locate the platform in the allotted time were gently guided there by the experimenter. Mice were allowed to stay on the platform for 15 s before being returned to their home cage between trials. Mice underwent training for 5 consecutive days. On MWM days 2-5, a probe trial was done prior to the day’s 4 training trials, in which the platform was removed from the maze, and animals were placed in the tank for 45 s to navigate the maze. Training ended after 5 days. A final probe trial was conducted 3 d later to test long-term memory for the trained location.

#### Repeated reversals water maze

Following the final 3 d probe trial in MWM, animals underwent three days of repeated reversal testing. Animals were given 8 training trials per day with a 15-20 min ITI to a novel location: SW, NW, and SE quadrants of the maze. Once training was completed for the specified location, the day began with a probe trial testing recall for the prior platform location. Following the probe trial, the platform was returned to the pool to again test in a novel quadrant.

#### Visible platform water maze

Following the final 24 hr probe trial for repeated reversal learning, all cues were removed from the walls, and the platform was returned to the pool but carrying a striped pole to mark its location. The platform was moved semi-randomly to test each quadrant of the pool twice thus totaling 8 cue trials with a 15-20 min ITI. Animals were placed into the pool from semi-random starting positions and allowed 60 sec to arrive to the cued platform where they were allowed to rest for 15 seconds before being dried and placed back in the warmed home cage.

#### Radial arm water maze

Visible cues were returned to the walls of the room in the same positions used for MWM and RR. The RAWM was created by installing clear Plexiglas triangular inserts into the existing water maze pool (41 cm on each side x 50 cm high), which resulted in six open arms joined at the center. Each arm measured 20 cm wide x 34 cm long, and the water was maintained at a depth of 38 cm. The static platform was located 3 cm from the end of one arm and submerged 1 cm below the water’s surface. Mice were placed into a different arm at the beginning of each trial, with the order of starting positions pseudo randomly selected before training, such that no trial began in an arm adjacent to the previous start position. Mice were allowed 60 s to navigate the maze. If a mouse failed to locate the platform in the allotted time, it was gently guided there by the experimenter and allowed to remain on the platform for 15 s before being returned to its home cage between trials. Mice underwent 8 training trials in 1 day with a 15 min ITI. Animals underwent a single probe trial 3 days later for long-term memory assessment.

### Tissue harvest

For T186S only, mice were euthanized by CO_2_, one hemisphere snap frozen on dry ice and stored at -80°C until use. For KO x tau and KI x tau animals, mice were euthanized by sodium pentobarbital overdose and transcardially perfused with ice-cold PBS containing 0.5 mM EDTA upon completion of behavioral testing at approximately 9 mo of age. Brains were removed and hemisected along the midline. One hemisphere was snap frozen for biochemistry, and the other hemisphere was immersion fixed in 4% PFA at 4°C for 48 hrs. The hemi-sected brains were cryoprotected in 30% sucrose at 4°C until equilibrated and then frozen in dry ice for histology use.

### Western blotting

Frozen hemispheres were sonicated in 5 vol:wt of RIPA buffer (50 mM Tris, 150 mM NaCl, 5 mM EDTA, 0.5% sodium deoxycholate, 1% NP-40, 0.1% SDS, pH 8.0) containing protease (Roche, 05892970001) and phosphatase inhibitors (Roche, 04906837001). Homogenates were centrifuged at 15 krpm for 10 min at 4^°^C, the supernatant collected, and protein concentration measured by BCA (Fisher, 23227). 50 ug of protein was diluted with Laemmli buffer and loaded without heat denaturation onto a Criterion 10-20% tris-HCl gel (Bio-Rad, 3450043) run in tris-glycine SDS buffer. The gel was transferred to nitrocellulose using the Transblot system (Bio-Rad, 1704271), and then blocked for 1 hr at RT in 1x PBS containing 5% non-fat dry milk and 0.1% Tween-20, followed by overnight incubation at 4°C with primary antibody diluted in block (Rb anti-TMEM106B, 1:1,000, Sigma HPA058342 and Ck anti-GAPDH, 1:10,000, Millipore ab2303). The following day, membranes were washed in 1x PBS containing 0.1% Tween-20 before being incubated for 1 hr at RT in secondary antibody diluted in block (Dk anti-Rb IgG 800 CW, 1:5,000, LI-COR 926-32213 and Dk anti-Ck IgY 680RD, 1:10,000, LI-COR 926-68075). Blots were then washed with PBS/Tween and visualized using an Odyssey Fc Imaging System (LI-COR). TMEM106B quantification was done using Image Studio Lite software, normalized to GAPDH.

### RNA extraction

Frozen hemi-brains were used to isolate RNA using a PureLink RNA Mini Kit (Invitrogen) according to the manufacturer’s directions (n=5 WT, 7 KO, 5 tau, and 7 KO:tau 3 males/group; n=12 WT, 12 KI, 12 tau, and 12 KI:tau, 6 males/group). In brief, brain hemispheres were homogenized in freshly prepared lysis buffer (0.6 ml per 30 mg of brain tissue) containing 1% *β*-mercaptoethanol using a TissueRuptor homogenizer (Qiagen).

Homogenates were centrifuged at 2,600 x g for 5 min at RT and supernatant was transferred to a clean RNase-free tube. One volume of 70% ethanol was added to the brain homogenate and 700 μl of the sample was transferred to a spin cartridge for RNA binding. This cartridge was centrifuged at 12,000 x g for 15 sec and then treated with DNase (On-Column PureLink DNase, Invitrogen) to obtain a DNA-free RNA sample. After several sequential washes, RNA was eluted with 50 μl of RNase-free water. Purified RNA yield and quality was assessed using a Nanodrop spectrophotometer and stored at -80°C until use.

### Transcriptomics

Library preparation, sequencing, read quality control, and mapping was performed by Novogene Co., LTD (Beijing, China). Briefly, library construction began by purifying mRNA that was then fragmented. cDNA was then synthesized from random hexamer primers and dTTP. cDNA underwent end repair, A-tailing, adapter ligation, size selection, amplification, and purification. RNA sequencing was performed by on the Illumina platform using paired end 150 bp mRNA sequencing with an expected read depth of 20M per sample. Data was filtered by removing adapter contamination, reads with uncertain nucleotide percentage above 10%, and reads with low quality scores constituting more than 50% of the read. Reads were mapped using HISAT2 v2.0.5 (Mortazavi et al., 2008) to mouse genome build mm10. The KO and KI analysis used separate samples arising from each colony and each contained all relevant experimental groups. Samples used in the KI analysis were generated from two different library preparations, and therefore counts from these samples were first normalized for batch effects using the Combat-Seq function in the sva package v3.44.0 with no covariates (Zhang et al., 2020). Samples from all experimental conditions and both sexes within the KI analysis were distributed evenly between the two batches. Differential gene expression analysis was performed using the edgeR pipeline v3.38.4 (Robinson et al., 2010). Differentially expressed genes from the KO and KI colonies were generated separately. Transcripts were first filtered to include only protein coding genes, and to further require cpm above 1 in 20% of any experimental group. 17,093 genes were included in analysis for the KO colony and 17,136 for the KI colony. Counts were then normalized to library size using the upper quartile method. Principle component analysis was first performed on normalized expression data to identify potential outlier samples. No samples were removed from downstream analysis. Results were calculated using the likelihood ratio testing parameter and Benjamin-Hochberg p-adjustment method. The RNA-seq data in this study was deposited at Gene Expression Omnibus (GEO) (http://www.ncbi.nlm.nih.gov/geo/) under accession ID GSE223376.

### Functional enrichment analysis

To perform functional enrichment of the KO:tau vs tau mice, we input the top DEGs (FDR<0.05) into STRINGdb (version 11.5) and applied Markov clustering followed by functional enrichment analysis of each cluster. For KI:tau vs tau transcriptional analysis, FDR<0.05 was too stringent, therefore we lowered the threshold to p<0.01. Functional analysis was performed in two different ways. One approach used STRINGdb (version 11.5) and applied Markov clustering followed by functional enrichment analysis of each cluster. Alternatively, the STRING network was entered into Cytoscape and we used the Hidef-Louvain clustering algorithm within the Community Detection Cytoscape application (V1.12.0) to generate clusters of DEGs using edge interaction strength as the weight column. Functional enrichment of each cluster was calculated using three different databases (gProfiler, iQuery and Enrichr). Networks were analyzed and represented using Cytoscape version: 3.9.1.

Gene set enrichment analysis (GSEA) was performed using GSEA v4.3.2 (Mootha et al., 2003; Subramanian et al., 2005) using default settings. Raw counts from protein coding genes that were normalized using upper quartile normalization were provided as input. For AD-specific GSEA, the KEGG AD pathway was run on its own (v2022, hsa05010). For subsequent functional analysis, we used the full KEGG pathway database. FDR q-value < 0.05 was considered significant.

### Integration of mouse DEGs with human AD DEGs

We used human brain RNAseq data from the AMP-AD knowledge portal to integrate DEGs identified in mice with transcriptionally dysregulated genes in AD (Allen et al., 2016; De Jager et al., 2018; Mostafavi et al., 2018). The AMP-AD data used for this comparison had been re-analyzed to normalize across different studies as detailed elsewhere, with definitions for AD patients and controls identical to those defined by Wan et al. (Wan et al., 2020). Specifically, we used datasets syn8484987, syn8466812, syn8456629 for the brain region-specific differential expression, using the data from AD vs controls only. The human brain samples were from ROSMAP (DLPFC 155 AD/ 86 Control), Mayo RNAseq (TCX 80-AD / 73-control) and Mount Sinai Brain Bank (FP 167-AD / 93-control, IFG 151-AD / 79-control, PHG 143-AD / 82-control, STG 151-AD / 89-control). P values for differential expression were adjusted for multiple hypothesis testing using false discovery rate estimation (FDR), and we selected genes with an adjusted p<0.05 based on the analysis in Wan et al. (Wan et al., 2020). For mouse DEGs, a cutoff of FDR<0.05 was used for KO:tau vs WT, tau vs WT, KO:tau vs tau, KI:tau vs WT, and tau vs WT, and a cutoff of p<0.01 for KI:tau vs tau. Significance of the overlap between human AD genes and mouse KO:tau DEGs (vs. tau alone) was calculated using Fisher’s exact test.

Correlation between mouse and human brain expression was calculated using Pearson’s correlation coefficient and significant differences between the regressions were calculated by ANOVA in full factorial modelling using standard least squares.

### Histology

Most histology for this study was performed by NeuroScience Associates using their MultiBrain technology (n=10 tau and 10 KO:tau, 5 males/group; n=10 tau and 10 KI:tau, 6 males/group). Frozen hemibrains were thawed and embedded into a gelatin matrix that was sectioned in the coronal plane at 35 um thickness. Sections were stored in cryoprotect buffer at -20 until use. A 1 in 12 series of free-floating sections was immunostained for AT8 or Iba1 using standard protocols and then lightly counterstained with hematoxylin (AT8 only) before mounting onto subbed slides, dehydrating through alcohols, and cover slipping with Permount. A 1 in 6 series of sections was used for Campbell-Switzer staining according to the protocol available at: https://www.neuroscienceassociates.com/reference/papers/alzheimers-disease-pathology-silver-stain/.

Additional MultiBrain sections from a 1 in 12 series were immunostained at BCM for CD-68 After quenching endogenous peroxidase with hydrogen peroxide, sections were blocked in serum and then incubated in primary antibody for 2 hr at RT (CD-68: 1:500, Bio-Rad clone FA-11, MCA1957GA), and then processed for secondary and HRP reagent using a Vectastain ABC kit according to the manufacturer’s instructions (Rat IgG, Vector Laboratories, PK-4004). Antibody binding was visualized with DAB-nickel (VWR, 101098-434). Sections were counterstained with nuclear fast red, mounted, dehydrated, and cover slipped with Permount.

Gallyas staining was done at BCM on a 1 in 12 series of MultiBrain sections. All glassware was acid-cleaned before use. Mounted tissue was bound to the slide by 5 min in 95% EtOH/37% formalin. Slides were rehydrated and then incubated at RT sequentially in potassium permanganate, oxalic acid, and periodic acid, each followed by a water rinse. Slides were then moved to silver iodide solution followed by acetic acid, before the stain was detected with Gallyas working solution. Slides were counterstained with nuclear fast red, dehydrated, and cover slipped with Permount.

### Histological quantitation

AT8, Iba1, and Campbell-Switzer-stained slides were imaged at NeuroScience Associates and uploaded into the Proscia Concentriq web-based digital pathology platform. CD68 slides were imaged using a Zeiss Axio Scan.Z1 at 10x magnification (Carl Zeiss AG, Oberkochen, Germany). For AT8 and Iba1 area analyses, 2 plane matched sections containing the dorsal hippocampus and spanning approximately -1.5 to -2.1 mm from bregma (tau and KO:tau), or 4-5 sections spanning approximately -1.5 to -2.8 mm (tau and KI:tau), were used for analysis. For AT8, Iba1, and CS silver, images were screen-captured from the Concentriq website at 3x magnification, saved as png files, imported into ImageJ v1.53s (NIH) as png files that were converted to 8-bit greyscale. Default (AT8) or Yen (Iba1) threshold was applied, then the maximum value was manually adjusted to better match the stain before calculating % area. For CD68 staining, six tif files per mouse, spanning approximately -1.82 to -4.24 mm from bregma, were imported into ImageJ, and quantification performed as for Iba1 using the Yen threshold and manually adjusting to match the stain before calculating area. Values were averaged across sections to calculate % area for each animal.

For volume measurements, a 1 in 6 series of Campbell-Switzer silver-stained sections was used for analysis, spanning the full rostro-caudal extent of the hippocampus; cortex and lateral ventricle were measured from the same sections (n=19-21 sections/mouse). Images were screen captured from the Concentriq website at 1.5x magnification and imported into ImageJ as png files. Regions of interest were manually traced, measured, and summed to calculate total volume. Summed values were then converted from pixels^2^ to mm^2^.

### Statistical analyses - behavior and histology

Statistical analyses were performed using GraphPad Prism 9. Comparison of two groups was done using unpaired two-tailed Student’s t-test or Mann-Whitney for non-parametric distribution. Comparison of three or more groups, or of two groups across days, was done using one- or two-way ANOVA followed by Tukey or Bonferroni post-hoc testing. Comparison of three or more groups with non-parametric distributions was done by Kruskal-Wallis followed by Dunn post-hoc testing. Survival curves were analyzed by Mantel-Cox log-rank test.

Statistical outliers were identified using either Grubb’s or ROUT testing in Prism. In only two cases were 2 animals removed from the same genotype (OF: 2 tau from the KI colony, and lateral ventricle area: 2 KI:tau). In all other cases, no more than 1 animal was removed as an outlier per genotype. One KO:tau animal that was impaired in visible platform testing was eliminated from all behavioral analyses.

## Acknowledgements

This work was supported by NIH grants R21AG056028, RF1AG058188, and P01AG066606-01A1 and -S1 (JLJ), Alzheimer’s Association grant ZEN-19-591129 (JLJ), F31AG067676-01A1 (CAW), T32NS043124 (support for GAE), Federal Work Study funds to BCM (support for QN and PJK), and by R01AG074009 (IAR and JLJ). This work was supported by several advanced technology core laboratories at BCM, including the RNA In Situ Hybridization Core, Bioengineering Core, and Genetically Engineered Rodent Models Core. BCM core laboratories were funded in part by NIH grants S10 OD016167 and P50 HD103555 (RNA ISH), P30 CA125123 (GERM), and P30 EY002520 (Bioeng). TMEM106B knockout germplasm was provided by the KOMP Repository and the Mouse Biology Program at the University of California Davis from ES cells generated by the Wellcome Trust Sanger Institute.

The results published here are in whole or in part based on data obtained from the AD Knowledge Portal (https://adknowledgeportal.org). The specific human AD RNAseq datasets used here were generated by: 1) the Rush Alzheimer’s Disease Center, Rush University Medical Center, Chicago. 2) Dr. Allan Levey from Emory University based on postmortem brain tissue collected by Dr. Eric Schadt from the Mount Sinai School of Medicine through the Mount Sinai VA Medical Center Brain Bank, and 3) the Mayo RNAseq study led by Dr. Nilüfer Ertekin-Taner, Mayo Clinic, Jacksonville, FL.

We thank Chelsea Jiaqi Zong, Zoe Lai, Jennifer Saldana, and Melissa Comstock for support with the mouse colonies; Gabriella Perez for help with imaging, Cecilia Ljungberg for help with slide scanning, I-Chih Tan for equipment support and repair, and Denise Lanza, Lan Liao, and Jason Heaney for help with mouse creation and cryorecovery.

## Supplemental Figures

**Supplemental Figure 1.**
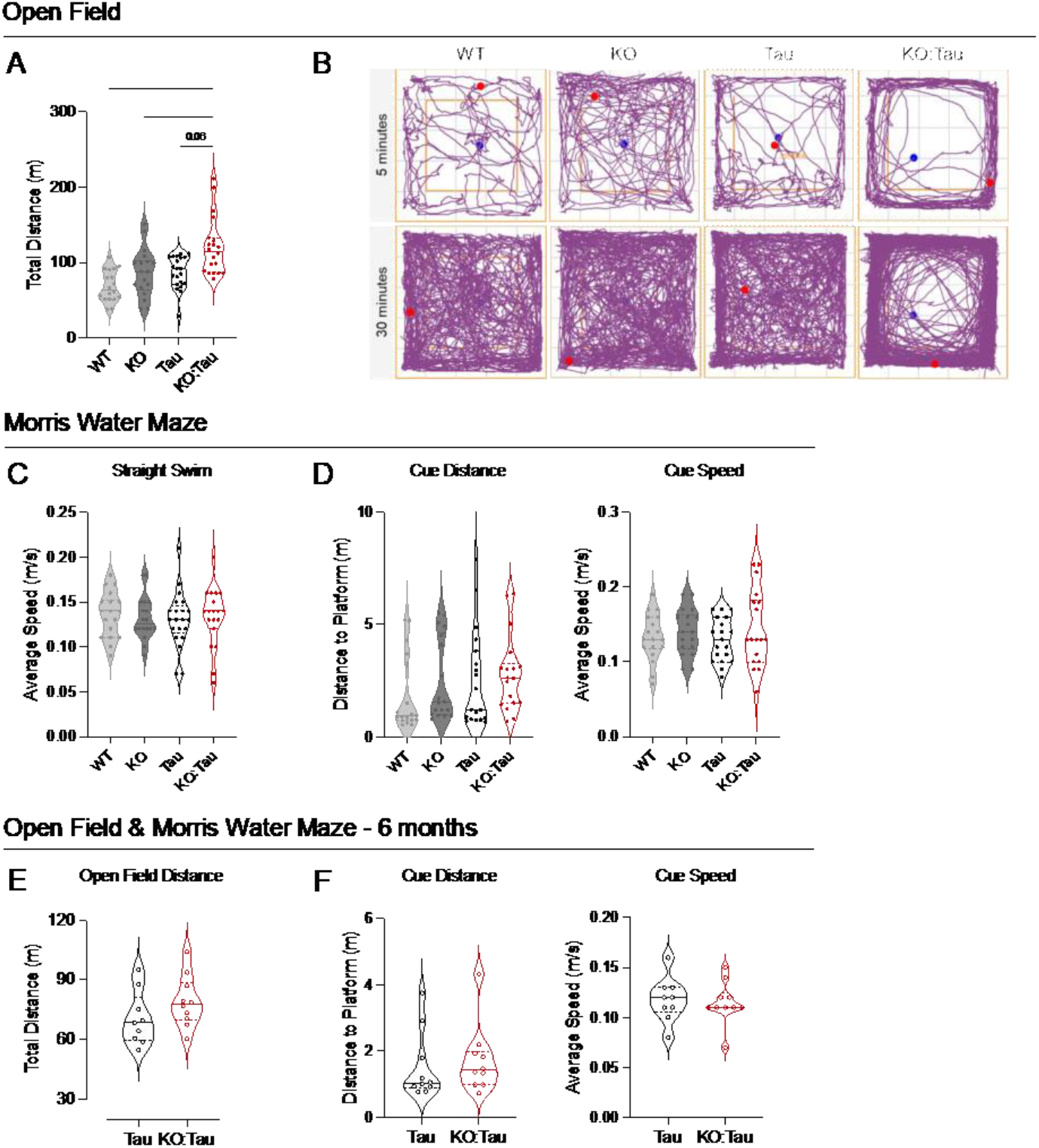
Basic assessment of locomotor and visual function uncovers hyperactivity in open field assay for tau mice lacking TMEM106B, but no other significant alterations. All animals were assessed for locomotor and visual function before and during cognitive testing using both open field and water-maze based measures. Data shown for panels A-D are from the 8 mo testing cohort; panels E-F are from the 6 mo cohort. A. Open field testing was conducted at the beginning of the behavioral test battery and revealed that KO:tau mice display hyperactivity. Graph shows total distance traveled during the 30 min test session. B. Example track plots showing the trajectory of a single representative animal in the open field during the first 5 min of testing (upper panels) and overall (lower panels). KO:tau animals were more likely to be thigmotaxic in the open field assay than other genotypes. C. Swim speed in a straight swim channel was tested prior to MWM training. In water, KO:tau mice performed similarly to all other genotypes. D. The swim speed (right) distance traveled (left) to a marked escape platform were measured at the end of repeated reversal testing. Again, no locomotor differences between genotypes were seen in water. E. Open field testing at 6 mo of age found no differences in total distance traveled between genotypes, suggesting that hyperactivity and thigmotaxis emerged with age and disease progression. F. As with the older cohort, neither swim speed nor distance traveled to the marked platform differed between genotypes in the 6 mo animals. * p<0.05, **** p<0.0001. See Table S1 for ANOVA statistics.

**Supplemental Figure 2.**
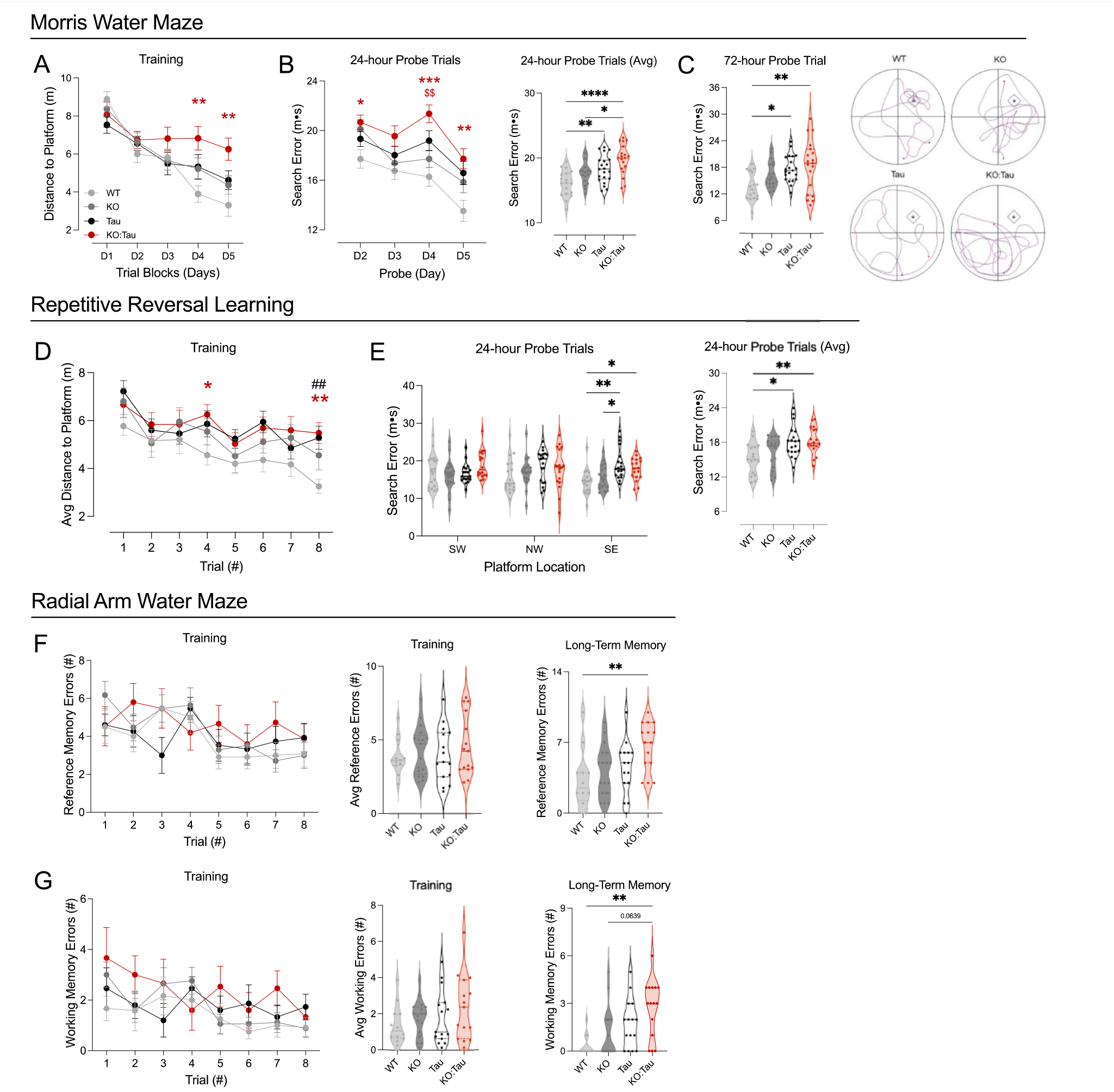
Loss of TMEM106B has limited impact on cognitive impairment in 9 mo old tau mice. A. Spatial learning was assessed using a standard Morris water maze which measured the distance to reach a hidden escape platform as a measure of learning. KO:tau mice performed significantly worse than WT on days 4 and 5, but the apparent difference in spatial learning between KO:tau and tau did not reach significance. B. Short-term memory for the escape location was tested by probe trials with the platform removed at the start of each training day 2-5. KO:tau mice performed significantly worse than WT and KO, but did not differ from tau on any individual testing day (left) or when averaged across testing days (right). C. Long-term memory for the escape location was tested 72 hours after the final training session. Again, there was no difference between KO:tau and tau mice, but both tau groups performed worse that WT (left). Track plots show example search paths for one animal of each genotype during the long-term probe trial (right). D. On completion of the MWM testing, cognitive flexibility was tested by moving the escape location each day for 3 consecutive days in the repeated reversal task. Distance to reach the escape location was measured over 8 trials/day, averaged across 3 training days. Both tau genotypes performed worse than WT, but there was no difference between KO:tau and tau in learning new locations at this age. E. Short-term memory for the changing platform location was tested 24 hr after training. Performance did not differ between KO:tau and tau for any given location (left), nor overall when averaged across the three locations, although both tau genotypes were worse than WT (right). F-G. The final test of the behavioral battery looked at both spatial reference memory and working memory using the radial arm water maze which converts the pool to a 6-arm format; performance is measured using incorrect arm entries (reference memory, F) or repeated arm entries (working memory, G). There was no difference in memory performance between KO:tau and tau during training, regardless of whether this was examined per trial (left) or averaged across trials (center). Long-term memory for the trained location differed only for KO:tau vs WT (right). * p<0.05, ** p<0.01, *** p<0.001, **** p<0.0001; ## p<0.01 tau vs WT, * p<0.05 KO:tau vs WT, ** p<0.01 KO:tau vs WT, *** p<0.001 KO:tau vs WT, $ p<0.05 KO:tau vs KO. See Table S1 for ANOVA statistics.

**Supplemental Figure 3.**
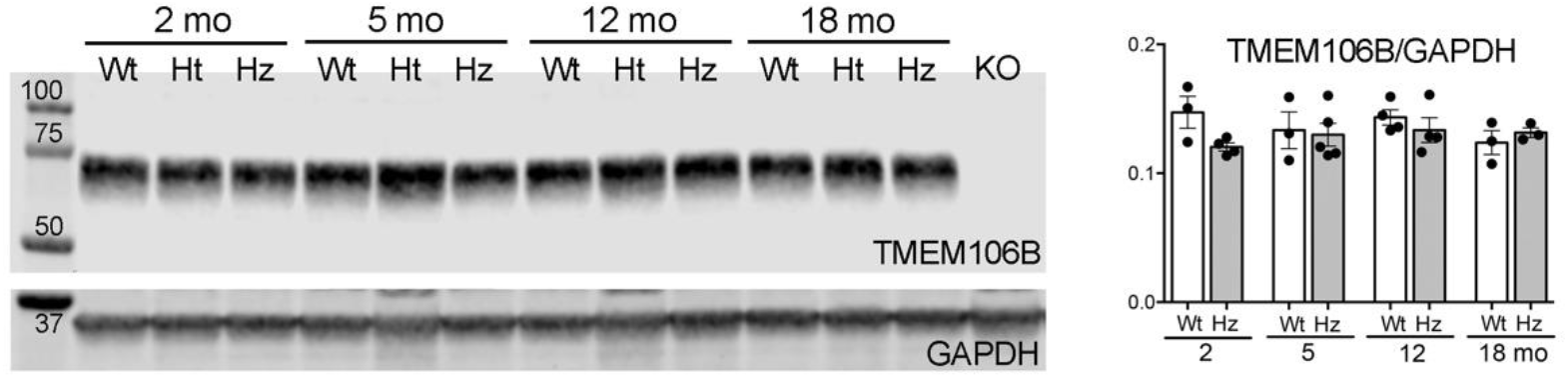
TMEM106B protein levels are unchanged by introduction of the T186S variant. Hemibrain homogenates were probed by Western blot to measure TMEM106B levels at 2, 5, 12, and 18 mo of age. No genotype differences were detected at any age. Wt: wild-type, Ht: heterozygous T186S, Hz: homozygous T186S. Final lane is brain extract from a TMEM106B KO animal.

**Supplemental Figure 4.**
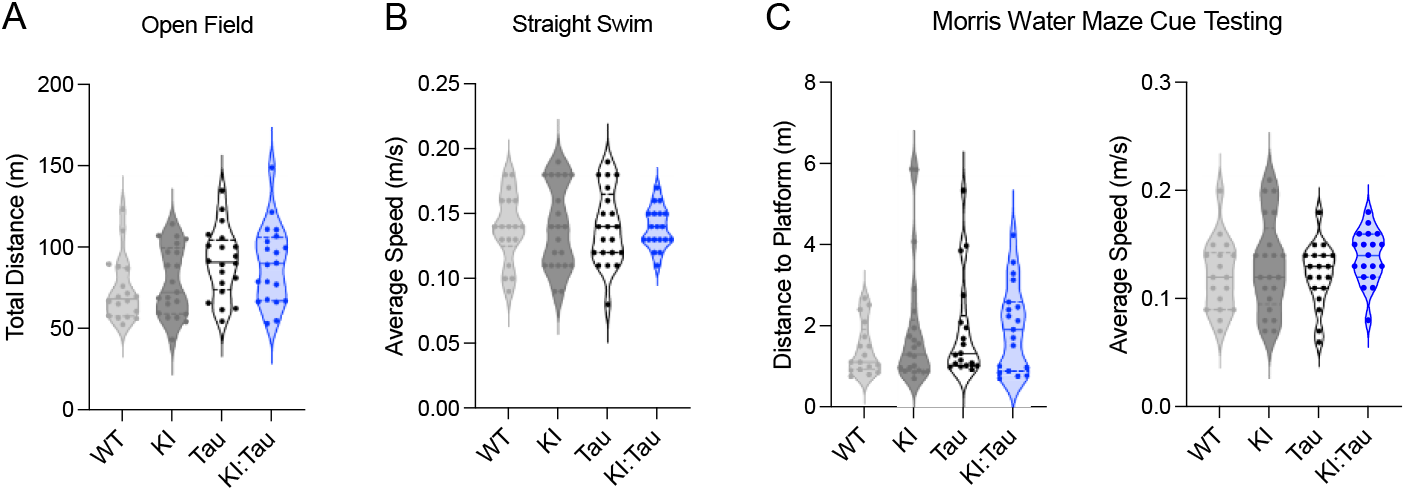
All genotypes of the KI x tau cohort display similar locomotor and visual ability. A. All genotypes traveled a similar distance during open field testing prior to cognitive testing. B. Swim speed in a straight channel tested prior to MWM training revealed no differences between genotypes. C. Both the distance traveled (left) and swim speed to reach a marked escape platform were similar between groups at the end of repeated reversal testing. See Table S1 for ANOVA statistics.

**Supplemental Figure 5.**
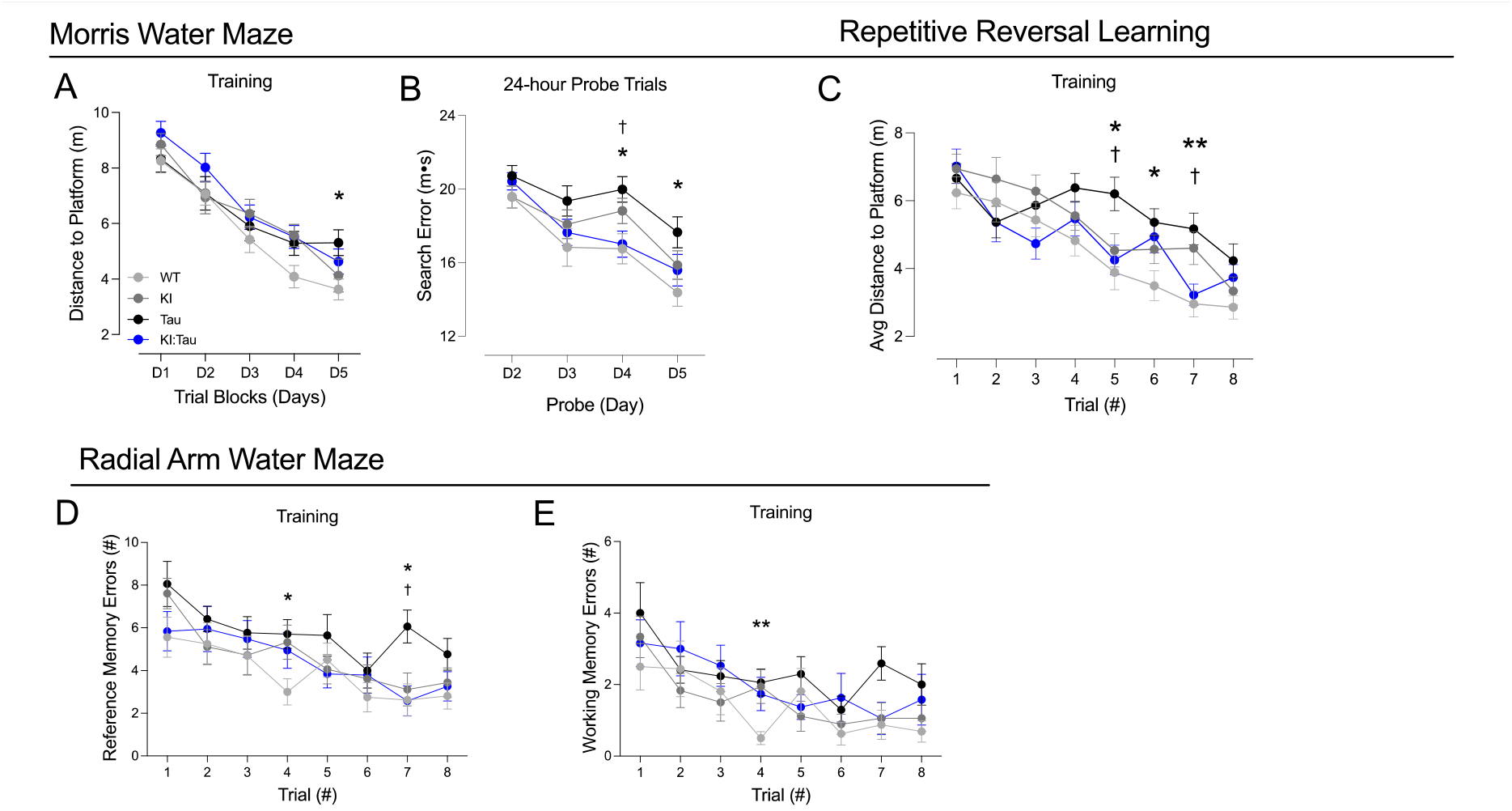
Full dataset for T186S x Tau cognitive testing. A. Morris water maze escape distance. Tau mice performed significantly worse than WT on day 5. B. MWM probe trials. Tau mice performed significantly worse than WT on days 4 and 5, and worse than KI:tau on day 4. C. Repeated reversal training. Tau mice performed worse than WT on trials 5-7, and worse than KI:tau on trials 5 and 7. D-E. Radial arm water maze performance: reference memory errors, D, and working memory errors, E, before reaching the escape platform. Tau mice performed worse than WT in reference memory on trials 4 and 7 and in working memory on trial 4. Tau mice were also worse than KI:tau in reference memory on trial 7. * p<0.05 tau vs WT, ** p<0.01 tau vs WT, † p<0.05 KI:tau vs tau. See Table 1 for complete ANOVA statistics.

**Supplemental Figure 6.**
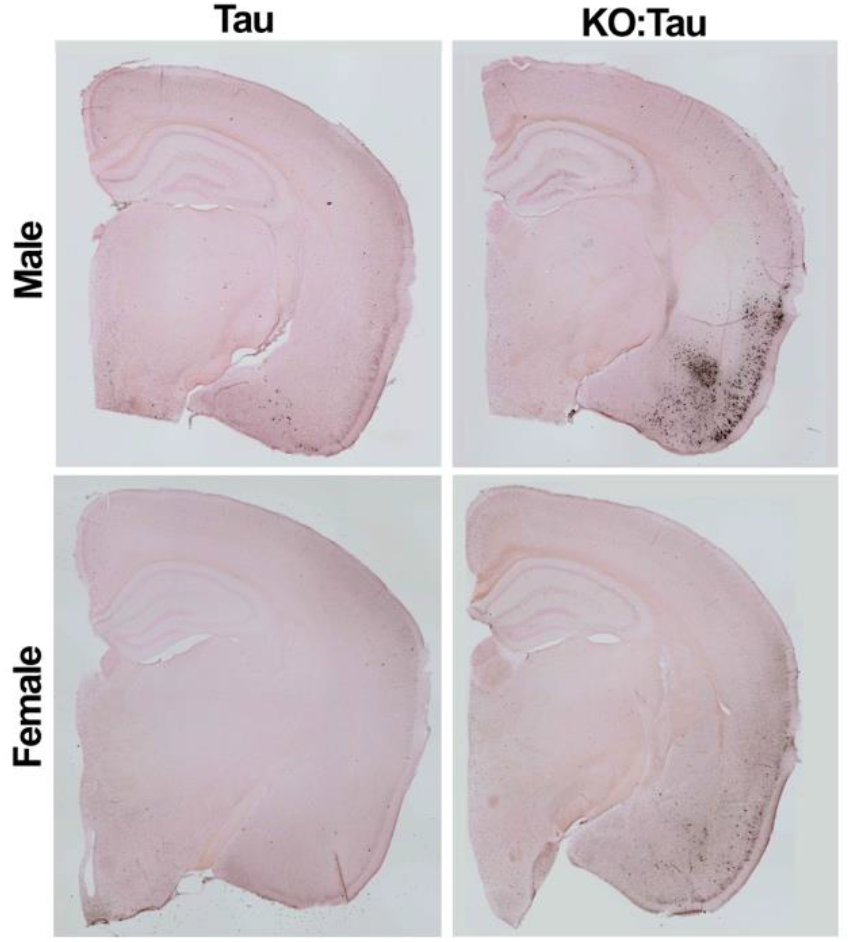
Neurofibrillary tangles were present by 6 mo of age in tau mice lacking TMEM106B. Gallyas staining was used to detect neurofibrillary tangles in tau and KO:tau mice harvested after behavioral testing at 6 mo of age. A few sparse tangles were found in male tau animals at this age, although rarely in females. In contrast, KO:tau mice had nearly all developed at least some tangles in the piriform area and/or amygdalar nuclei by this age across both sexes.

**Supplemental Figure 7.**
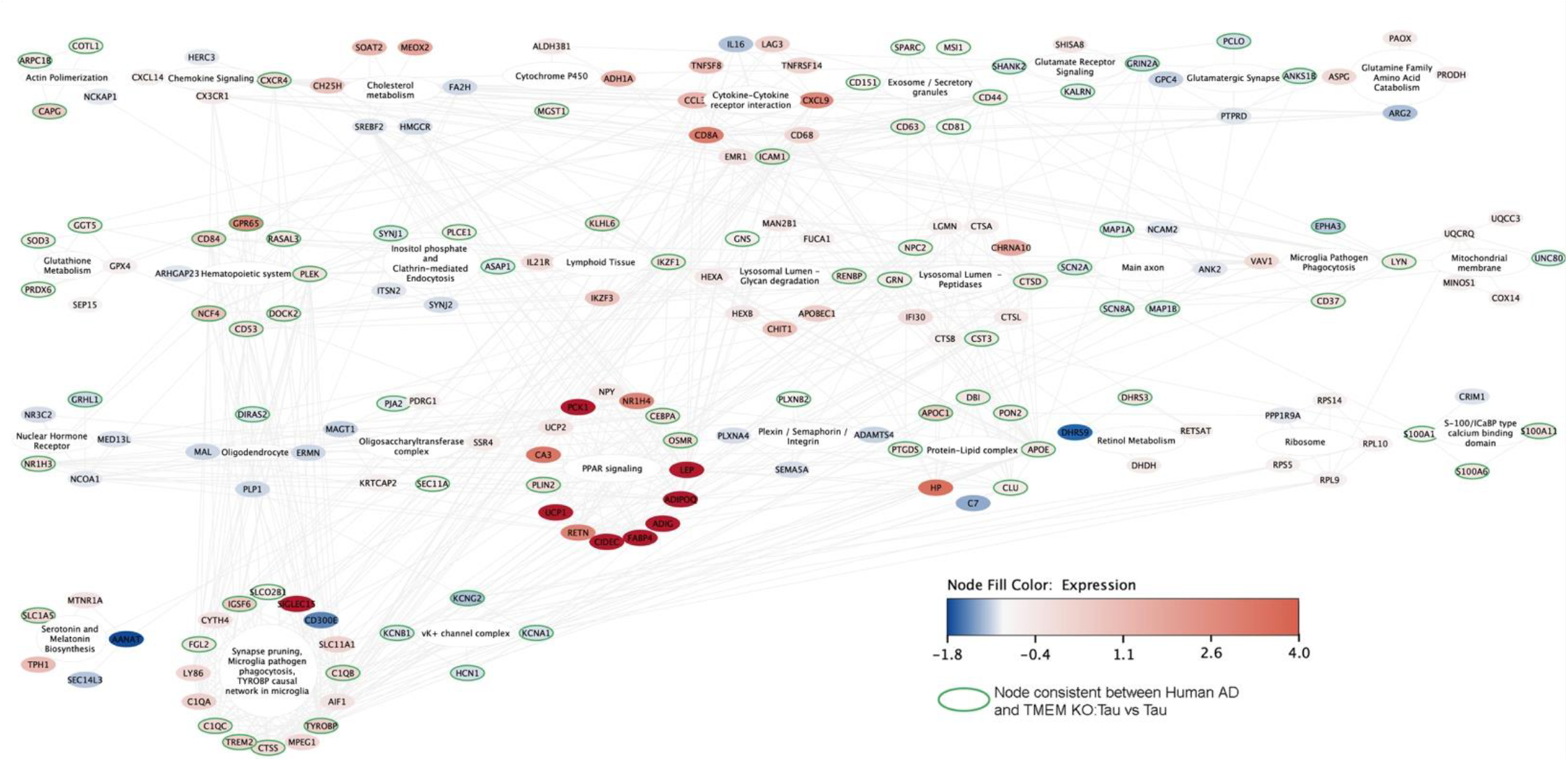
Functional enrichment analysis for genes identified from comparison of KO:tau vs tau. Log2 fold-change between genotypes is indicated by the heat map coloring for each gene. DEGs that were concordant between human AD and mouse are outlined in green. Edges connecting nodes within different modules are indicated in grey. The functionally enriched pathways are labeled inside each module. The TYROBP module showed a high degree of interconnectedness with multiple other modules. PPAR signaling is also notable for the number of genes and the degree of change upon TMEM106B deletion. The network was built using STRINGdb v11.5 and visualized using Cytoscape.

**Supplemental Figure 8.**
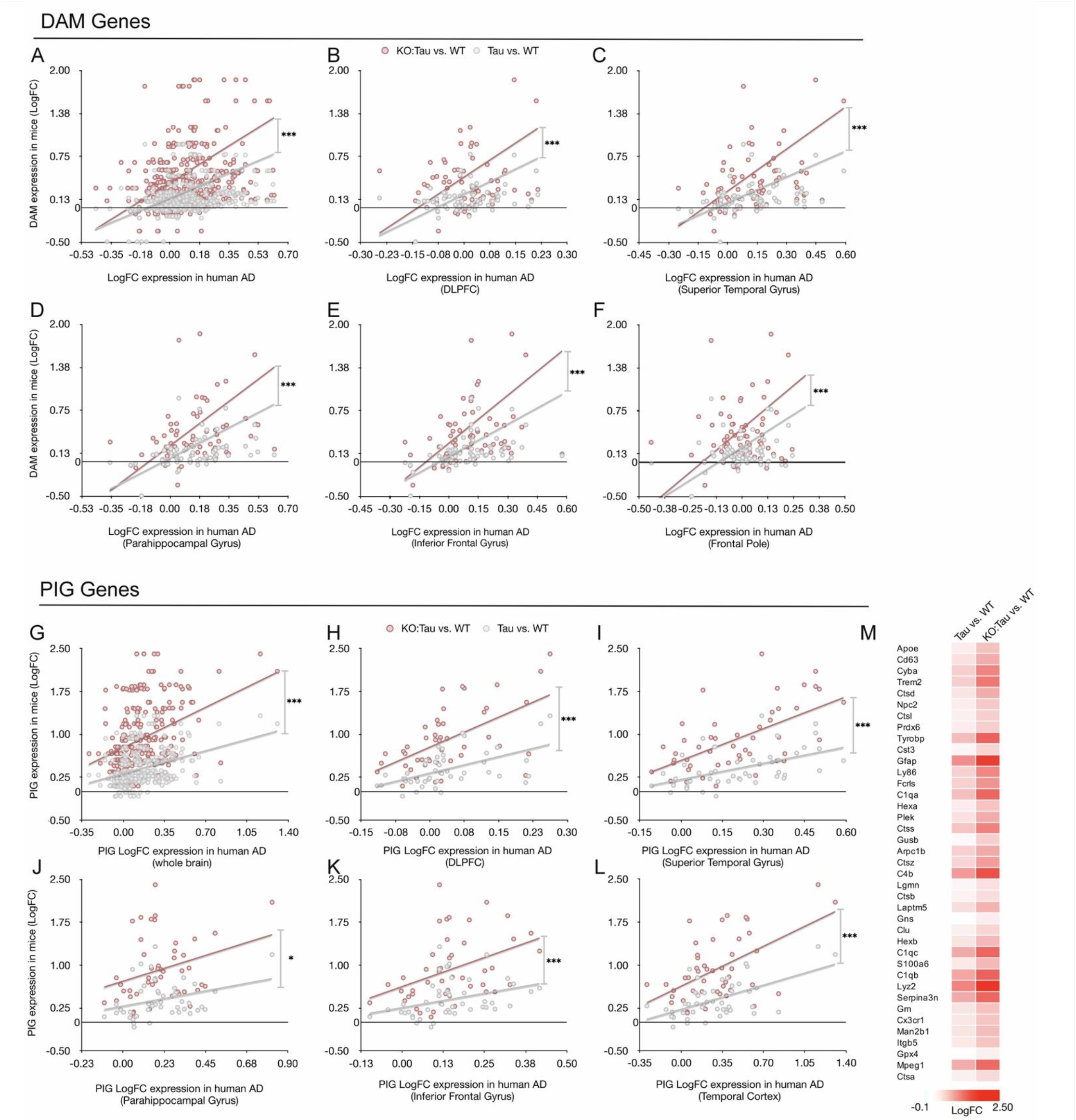
Correlation between mouse and human DAM and PIG gene expression. DAM genes were identified from Keren-Shaul et al. (Keren-Shaul et al., 2017); PIG genes were identified from Chen et al. (Chen et al., 2020). Log2 fold change for KO:tau vs WT (pink) and tau vs WT (grey) are plotted as a function of log2 FC for human AD vs HC. A-F. Correlation between human and mouse gene expression for DAM genes. G-L. Correlation between human and mouse gene expression for PIG genes. A and G. Combined values for human vs mouse correlations including all 5 human brain regions examined. B-F and H-L. Correlations for human vs mouse plotted separately for each human brain region. DLPFC=dorsolateral prefrontal cortex. M. Heat map showing the relative expression of PIG genes that were differentially expressed between KO:tau and tau. *p<0.05, *** p<0.001

**Supplemental Figure 9.**
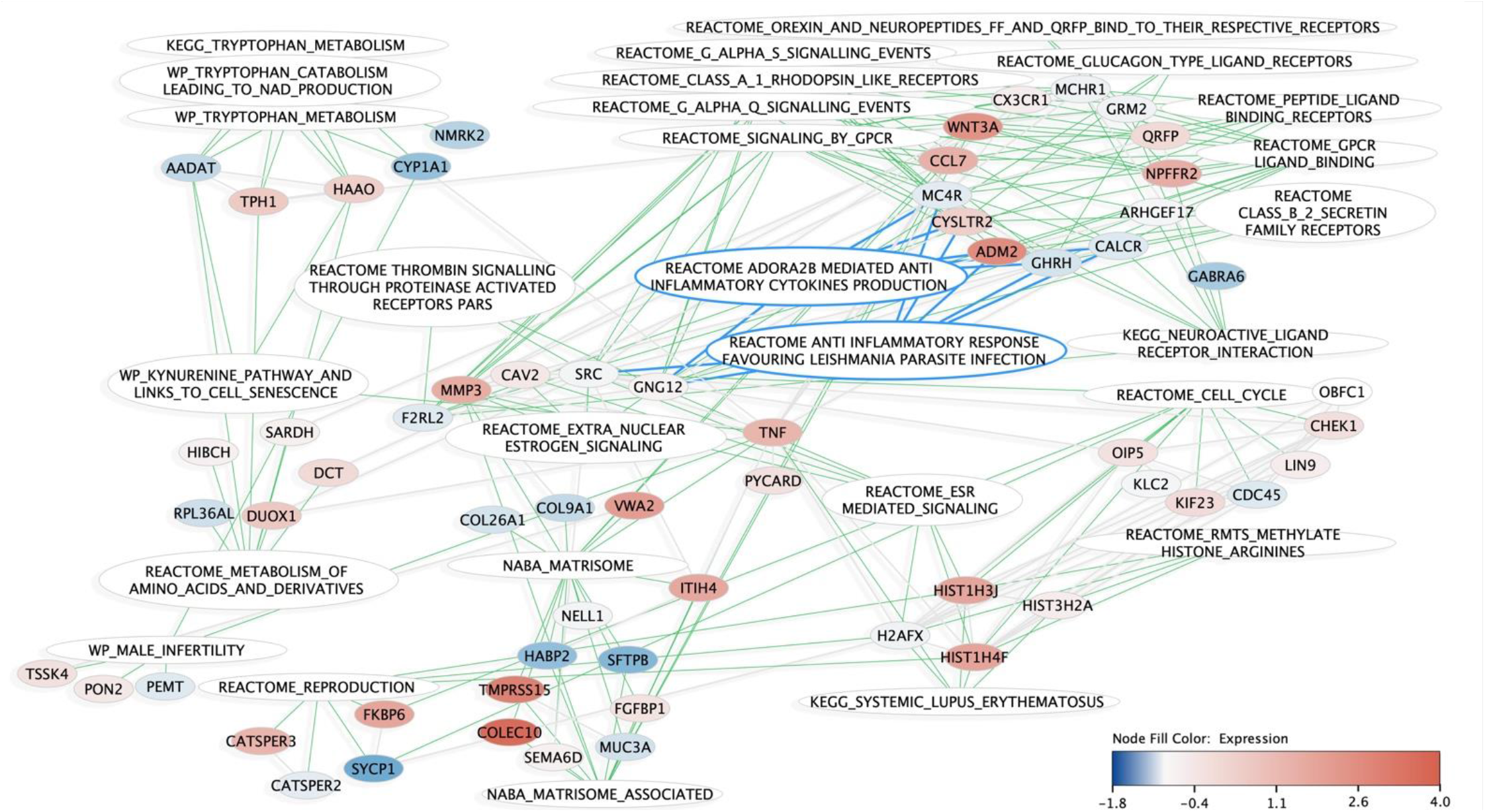
Functional enrichment analysis for genes identified from comparison of KI:tau vs tau. Log2 fold-change between genotypes is indicated by the heat map coloring for each gene. Green edges connect genes to the modules they are involved in, blue edges highlight connections and pathways that inhibit inflammation. The network was built using STRINGdb and functional enrichment was calculated using MSigDB - GSEA and visualized using Cytoscape.

## References

Allen, M., Carrasquillo, M.M., Funk, C., Heavner, B.D., Zou, F., Younkin, C.S., Burgess, J.D., Chai, H.S., Crook, J., Eddy, J.A., et al. (2016). Human whole genome genotype and transcriptome data for Alzheimer’s and other neurodegenerative diseases. Sci Data 3, 160089.

Bennett, D.A., Schneider, J.A., Arvanitakis, Z., and Wilson, R.S. (2012). Overview and findings from the religious orders study. Curr Alzheimer Res 9, 628–645.

Briggs, D.I., Defensor, E., Memar Ardestani, P., Yi, B., Halpain, M., Seabrook, G., and Shamloo, M. (2017). Role of Endoplasmic Reticulum Stress in Learning and Memory Impairment and Alzheimer’s Disease-Like Neuropathology in the PS19 and APP(Swe) Mouse Models of Tauopathy and Amyloidosis. eNeuro 4.

Burmeister, A.R., and Marriott, I. (2018). The Interleukin-10 Family of Cytokines and Their Role in the CNS. Frontiers in cellular neuroscience 12, 458.

Cabron, A.S., Borgmeyer, U., Richter, J., Peisker, H., Gutbrod, K., Dormann, P., Capell, A., and Damme, M. (2023). Lack of a protective effect of the Tmem106b “protective SNP” in the Grn knockout mouse model for frontotemporal lobar degeneration. Acta Neuropathol Commun 11, 21.

Cen, L.P., Ng, T.K., Chu, W.K., and Pang, C.P. (2022). Growth hormone-releasing hormone receptor signaling in experimental ocular inflammation and neuroprotection. Neural Regen Res 17, 2643–2648.

Chang, A., Xiang, X., Wang, J., Lee, C., Arakhamia, T., Simjanoska, M., Wang, C., Carlomagno, Y., Zhang, G., Dhingra, S., et al. (2022). Homotypic fibrillization of TMEM106B across diverse neurodegenerative diseases. Cell 185, 1346–1355 e1315.

Chen, K., Fu, Q., Li, D., Wu, Y., Sun, S., and Zhang, X. (2016). Wnt3a suppresses Pseudomonas aeruginosa-induced inflammation and promotes bacterial killing in macrophages. Mol Med Rep 13, 2439–2446.

Chen, W.T., Lu, A., Craessaerts, K., Pavie, B., Sala Frigerio, C., Corthout, N., Qian, X., Lalakova, J., Kuhnemund, M., Voytyuk, I., et al. (2020). Spatial Transcriptomics and In Situ Sequencing to Study Alzheimer’s Disease. Cell 182, 976–991 e919.

Colonna, M. (2023). The biology of TREM receptors. Nat Rev Immunol, 1–15.

Davies, J., Chen, J., Pink, R., Carter, D., Saunders, N., Sotiriadis, G., Bai, B., Pan, Y., Howlett, D., Payne, A., et al. (2015). Orexin receptors exert a neuroprotective effect in Alzheimer’s disease (AD) via heterodimerization with GPR103. Sci Rep 5, 12584.

De Jager, P.L., Ma, Y., McCabe, C., Xu, J., Vardarajan, B.N., Felsky, D., Klein, H.U., White, C.C., Peters, M.A., Lodgson, B., et al. (2018). A multi-omic atlas of the human frontal cortex for aging and Alzheimer’s disease research. Sci Data 5, 180142.

Feng, T., Lacrampe, A., and Hu, F. (2021). Physiological and pathological functions of TMEM106B: a gene associated with brain aging and multiple brain disorders. Acta Neuropathol 141, 327–339.

Feng, T., Mai, S., Roscoe, J.M., Sheng, R.R., Ullah, M., Zhang, J., Katz, II, Yu, H., Xiong, W., and Hu, F. (2020). Loss of TMEM106B and PGRN leads to severe lysosomal abnormalities and neurodegeneration in mice. EMBO Rep 21, e50219.

Fowler, S.W., Chiang, A.C., Savjani, R.R., Larson, M.E., Sherman, M.A., Schuler, D.R., Cirrito, J.R., Lesne, S.E., and Jankowsky, J.L. (2014). Genetic modulation of soluble abeta rescues cognitive and synaptic impairment in a mouse model of Alzheimer’s disease. J Neurosci 34, 7871–7885.

Gallagher, M.D., Posavi, M., Huang, P., Unger, T.L., Berlyand, Y., Gruenewald, A.L., Chesi, A., Manduchi, E., Wells, A.D., Grant, S.F.A., et al. (2017). A Dementia-Associated Risk Variant near TMEM106B Alters Chromatin Architecture and Gene Expression. American journal of human genetics 101, 643–663.

Heuberger, D.M., and Schuepbach, R.A. (2019). Protease-activated receptors (PARs): mechanisms of action and potential therapeutic modulators in PAR-driven inflammatory diseases. Thromb J 17, 4.

Jiang, Y.X., Cao, Q., Sawaya, M.R., Abskharon, R., Ge, P., DeTure, M., Dickson, D.W., Fu, J.Y., Ogorzalek Loo, R.R., Loo, J.A., and Eisenberg, D.S. (2022). Amyloid fibrils in FTLD-TDP are composed of TMEM106B and not TDP-43. Nature 605, 304–309.

Jun, M.H., Han, J.H., Lee, Y.K., Jang, D.J., Kaang, B.K., and Lee, J.A. (2015). TMEM106B, a frontotemporal lobar dementia (FTLD) modifier, associates with FTD-3-linked CHMP2B, a complex of ESCRT-III. Mol Brain 8, 85.

Keren-Shaul, H., Spinrad, A., Weiner, A., Matcovitch-Natan, O., Dvir-Szternfeld, R., Ulland, T.K., David, E., Baruch, K., Lara-Astaiso, D., Toth, B., et al. (2017). A Unique Microglia Type Associated with Restricting Development of Alzheimer’s Disease. Cell 169, 1276–1290 e1217.

Klein, Z.A., Takahashi, H., Ma, M., Stagi, M., Zhou, M., Lam, T.T., and Strittmatter, S.M. (2017). Loss of TMEM106B Ameliorates Lysosomal and Frontotemporal Dementia-Related Phenotypes in Progranulin-Deficient Mice. Neuron 95, 281–296 e286.

Larson, K.C., Lipko, M., Dabrowski, M., and Draper, M.P. (2010). Gng12 is a novel negative regulator of LPS-induced inflammation in the microglial cell line BV-2. Inflamm Res 59, 15–22.

Lewandoski, M., Meyers, E.N., and Martin, G.R. (1997). Analysis of Fgf8 gene function in vertebrate development. Cold Spring Harb Symp Quant Biol 62, 159–168.

Li, T., Wang, W., Gong, S., Sun, H., Zhang, H., Yang, A.G., Chen, Y.H., and Li, X. (2018). Genome-wide analysis reveals TNFAIP8L2 as an immune checkpoint regulator of inflammation and metabolism. Mol Immunol 99, 154–162.

Li, Z., Farias, F.H.G., Dube, U., Del-Aguila, J.L., Mihindukulasuriya, K.A., Fernandez, M.V., Ibanez, L., Budde, J.P., Wang, F., Lake, A.M., et al. (2020). The TMEM106B FTLD-protective variant, rs1990621, is also associated with increased neuronal proportion. Acta Neuropathol 139, 45–61.

Luningschror, P., Werner, G., Stroobants, S., Kakuta, S., Dombert, B., Sinske, D., Wanner, R., Lullmann-Rauch, R., Wefers, B., Wurst, W., et al. (2020). The FTLD Risk Factor TMEM106B Regulates the Transport of Lysosomes at the Axon Initial Segment of Motoneurons. Cell reports 30, 3506–3519 e3506.

Min, S.W., Chen, X., Tracy, T.E., Li, Y., Zhou, Y., Wang, C., Shirakawa, K., Minami, S.S., Defensor, E., Mok, S.A., et al. (2015). Critical role of acetylation in tau-mediated neurodegeneration and cognitive deficits. Nat Med 21, 1154–1162.

Mootha, V.K., Lindgren, C.M., Eriksson, K.F., Subramanian, A., Sihag, S., Lehar, J., Puigserver, P., Carlsson, E., Ridderstrale, M., Laurila, E., et al. (2003). PGC-1alpha-responsive genes involved in oxidative phosphorylation are coordinately downregulated in human diabetes. Nat Genet 34, 267–273.

Mortazavi, A., Williams, B.A., McCue, K., Schaeffer, L., and Wold, B. (2008). Mapping and quantifying mammalian transcriptomes by RNA-Seq. Nat Methods 5, 621–628.

Mostafavi, S., Gaiteri, C., Sullivan, S.E., White, C.C., Tasaki, S., Xu, J., Taga, M., Klein, H.U., Patrick, E., Komashko, V., et al. (2018). A molecular network of the aging human brain provides insights into the pathology and cognitive decline of Alzheimer’s disease. Nat Neurosci 21, 811–819.

Nicholson, A.M., Finch, N.A., Wojtas, A., Baker, M.C., Perkerson, R.B., 3rd, Castanedes-Casey, M., Rousseau, L., Benussi, L., Binetti, G., Ghidoni, R., et al. (2013). TMEM106B p.T185S regulates TMEM106B protein levels: implications for frontotemporal dementia. Journal of neurochemistry 126, 781–791.

Nicholson, A.M., and Rademakers, R. (2016). What we know about TMEM106B in neurodegeneration. Acta Neuropathol 132, 639–651.

Nicholson, A.M., Zhou, X., Perkerson, R.B., Parsons, T.M., Chew, J., Brooks, M., DeJesus-Hernandez, M., Finch, N.A., Matchett, B.J., Kurti, A., et al. (2018). Loss of Tmem106b is unable to ameliorate frontotemporal dementia-like phenotypes in an AAV mouse model of C9ORF72-repeat induced toxicity. Acta Neuropathol Commun 6, 42.

Perneel, J., and Rademakers, R. (2022). Identification of TMEM106B amyloid fibrils provides an updated view of TMEM106B biology in health and disease. Acta Neuropathol 144, 807–819.

Rhinn, H., and Abeliovich, A. (2017). Differential Aging Analysis in Human Cerebral Cortex Identifies Variants in TMEM106B and GRN that Regulate Aging Phenotypes. Cell Syst 4, 404–415 e405.

Robinson, M.D., McCarthy, D.J., and Smyth, G.K. (2010). edgeR: a Bioconductor package for differential expression analysis of digital gene expression data. Bioinformatics 26, 139–140.

Rodriguez, C.I., Buchholz, F., Galloway, J., Sequerra, R., Kasper, J., Ayala, R., Stewart, A.F., and Dymecki, S.M. (2000). High-efficiency deleter mice show that FLPe is an alternative to Cre-loxP. Nat Genet 25, 139–140.

Sadagurski, M., Landeryou, T., Cady, G., Kopchick, J.J., List, E.O., Berryman, D.E., Bartke, A., and Miller, R.A. (2015). Growth hormone modulates hypothalamic inflammation in long-lived pituitary dwarf mice. Aging Cell 14, 1045–1054.

Schweighauser, M., Arseni, D., Bacioglu, M., Huang, M., Lovestam, S., Shi, Y., Yang, Y., Zhang, W., Kotecha, A., Garringer, H.J., et al. (2022). Age-dependent formation of TMEM106B amyloid filaments in human brains. Nature 605, 310–314.

Subramanian, A., Tamayo, P., Mootha, V.K., Mukherjee, S., Ebert, B.L., Gillette, M.A., Paulovich, A., Pomeroy, S.L., Golub, T.R., Lander, E.S., and Mesirov, J.P. (2005). Gene set enrichment analysis: a knowledge-based approach for interpreting genome-wide expression profiles. Proc Natl Acad Sci U S A 102, 15545–15550.

Takeuchi, H., Iba, M., Inoue, H., Higuchi, M., Takao, K., Tsukita, K., Karatsu, Y., Iwamoto, Y., Miyakawa, T., Suhara, T., et al. (2011). P301S mutant human tau transgenic mice manifest early symptoms of human tauopathies with dementia and altered sensorimotor gating. PloS one 6, e21050.

Van Deerlin, V.M., Sleiman, P.M., Martinez-Lage, M., Chen-Plotkin, A., Wang, L.S., Graff-Radford, N.R., Dickson, D.W., Rademakers, R., Boeve, B.F., Grossman, M., et al. (2010). Common variants at 7p21 are associated with frontotemporal lobar degeneration with TDP-43 inclusions. Nat Genet 42, 234–239.

van der Touw, W., Chen, H.M., Pan, P.Y., and Chen, S.H. (2017). LILRB receptor-mediated regulation of myeloid cell maturation and function. Cancer Immunol Immunother 66, 1079–1087.

van der Zee, J., Van Langenhove, T., Kleinberger, G., Sleegers, K., Engelborghs, S., Vandenberghe, R., Santens, P., Van den Broeck, M., Joris, G., Brys, J., et al. (2011). TMEM106B is associated with frontotemporal lobar degeneration in a clinically diagnosed patient cohort. Brain 134, 808–815.

Wan, Y.W., Al-Ouran, R., Mangleburg, C.G., Perumal, T.M., Lee, T.V., Allison, K., Swarup, V., Funk, C.C., Gaiteri, C., Allen, M., et al. (2020). Meta-Analysis of the Alzheimer’s Disease Human Brain Transcriptome and Functional Dissection in Mouse Models. Cell reports 32, 107908.

Wang, M., Beckmann, N.D., Roussos, P., Wang, E., Zhou, X., Wang, Q., Ming, C., Neff, R., Ma, W., Fullard, J.F., et al. (2018). The Mount Sinai cohort of large-scale genomic, transcriptomic and proteomic data in Alzheimer’s disease. Sci Data 5, 180185.

Waqas, S.F.H., Hoang, A.C., Lin, Y.T., Ampem, G., Azegrouz, H., Balogh, L., Thuroczy, J., Chen, J.C., Gerling, I.C., Nam, S., et al. (2017). Neuropeptide FF increases M2 activation and self-renewal of adipose tissue macrophages. J Clin Invest 127, 2842–2854.

Werner, G., Damme, M., Schludi, M., Gnorich, J., Wind, K., Fellerer, K., Wefers, B., Wurst, W., Edbauer, D., Brendel, M., et al. (2020). Loss of TMEM106B potentiates lysosomal and FTLD-like pathology in progranulin-deficient mice. EMBO Rep 21, e50241.

White, C.C., Yang, H.S., Yu, L., Chibnik, L.B., Dawe, R.J., Yang, J., Klein, H.U., Felsky, D., Ramos-Miguel, A., Arfanakis, K., et al. (2017). Identification of genes associated with dissociation of cognitive performance and neuropathological burden: Multistep analysis of genetic, epigenetic, and transcriptional data. PLoS medicine 14, e1002287.

Yanamandra, K., Kfoury, N., Jiang, H., Mahan, T.E., Ma, S., Maloney, S.E., Wozniak, D.F., Diamond, M.I., and Holtzman, D.M. (2013). Anti-tau antibodies that block tau aggregate seeding in vitro markedly decrease pathology and improve cognition in vivo. Neuron 80, 402–414.

Yoshiyama, Y., Higuchi, M., Zhang, B., Huang, S.M., Iwata, N., Saido, T.C., Maeda, J., Suhara, T., Trojanowski, J.Q., and Lee, V.M. (2007). Synapse loss and microglial activation precede tangles in a P301S tauopathy mouse model. Neuron 53, 337–351.

Zhang, B., Carroll, J., Trojanowski, J.Q., Yao, Y., Iba, M., Potuzak, J.S., Hogan, A.M., Xie, S.X., Ballatore, C., Smith, A.B., 3rd, et al. (2012). The microtubule-stabilizing agent, epothilone D, reduces axonal dysfunction, neurotoxicity, cognitive deficits, and Alzheimer-like pathology in an interventional study with aged tau transgenic mice. J Neurosci 32, 3601–3611.

Zhang, D., Lu, Z., Man, J., Cui, K., Fu, X., Yu, L., Gao, Y., Liao, L., Xiao, Q., Guo, R., et al. (2019). Wnt-3a alleviates neuroinflammation after ischemic stroke by modulating the responses of microglia/macrophages and astrocytes. Int Immunopharmacol 75, 105760.

Zhang, Y., Parmigiani, G., and Johnson, W.E. (2020). ComBat-seq: batch effect adjustment for RNA-seq count data. NAR Genom Bioinform 2, lqaa078.

Zhou, X., Brooks, M., Jiang, P., Koga, S., Zuberi, A.R., Baker, M.C., Parsons, T.M., Castanedes-Casey, M., Phillips, V., Librero, A.L., et al. (2020a). Loss of Tmem106b exacerbates FTLD pathologies and causes motor deficits in progranulin-deficient mice. EMBO Rep 21, e50197.

Zhou, X., Nicholson, A.M., Ren, Y., Brooks, M., Jiang, P., Zuberi, A., Phuoc, H.N., Perkerson, R.B., Matchett, B., Parsons, T.M., et al. (2020b). Loss of TMEM106B leads to myelination deficits: implications for frontotemporal dementia treatment strategies. Brain 143, 1905–1919.

Zhu, X.C., Yu, J.T., Jiang, T., Wang, P., Cao, L., and Tan, L. (2015). CR1 in Alzheimer’s disease. Molecular neurobiology 51, 753–765.

Ziogas, D.C., Karagiannis, A.K., Geiger, B.M., Gras-Miralles, B., Najarian, R., Reizes, O., Fitzpatrick, L.R., and Kokkotou, E. (2014). Inflammation-induced functional connectivity of melanin-concentrating hormone and IL-10. Peptides 55, 58–64.

